# Quantitative analysis of DNA-GATA1 binding alterations linked to hematopoietic disorders

**DOI:** 10.1101/2024.08.26.609639

**Authors:** Boglarka Zambo, Bastien Morlet, Luc Negroni, Orsolya Mózner, Balázs Sarkadi, Gilles Travé, Gergo Gogl

## Abstract

GATA1 is a crucial transcription factor involved in hematopoiesis, and mutations in this gene are linked to severe hematological disorders, including anemia, thrombocytopenia, Down-syndrome related transient abnormal myelopoiesis (DS-TAM) and acute megakaryocytic leukemia (DS-AMKL). Despite significant clinical interest in the molecular level characterization of GATA1 mutations, a comprehensive understanding of their impact on DNA binding is limited. Efforts to conduct detailed studies on full-length recombinant GATA1 have faced significant technical challenges, while alternative approaches are limited by low throughput or qualitative nature. Here, we introduce a native holdup (nHU) assay designed to systematically quantify DNA-protein interactions and suitable for studying the impact of transcription factor mutations on DNA-binding affinity. First, we demonstrate, using the erythroid-specific *ATP2B4* promoter as a model, that nHU can capture sequence-specific interactions and detect even subtle differences in DNA binding affinities of GATA1. Then, we quantitatively characterize the impact of pathological mutations on DNA binding affinities in the context of full-length human GATA1. Our findings reveal that the short isoform, lacking the N-terminal transactivation domain (N-TAD), binds to DNA with increased affinity, while the R307C mutation reduces binding to the *ATP2B4* erythroid promoter. In harmony with these observations, GATA1s exhibits increased functional activity, while the R307C mutation results in decreased activity. This study demonstrates the power of the nHU assay for studying DNA interactions of transcription factor variants and providing insight into the molecular mechanism of related diseases.

## Introduction

The erythroid transcription factor (TF) GATA1 is an essential protein involved in the differentiation of myeloid progenitor cells to mature blood cells in the bone marrow (1, 2). Complete loss of the GATA1 in mice is lethal at embryonic stage due to the absence of red blood cells, and mutations in the gene can alter proper erythropoiesis (3–6). GATA1 is composed of two C4-type zinc fingers, often referred to as the N- and C-terminal fingers (NF and CF, respectively, Figure 1A) (7). The CF binds to DNA with high affinity and specificity through (A/T)GATA(A/G) sequences, while the NF displays weaker affinity to similar sites and appears to rather play a modulatory role by interacting with other proteins such as FOG1, or LMO2 (8–10). In the past decades, many mutations have been identified in patients suffering from severe anemia, thrombocytopenia and/or β-thalassemia (Figure 1A). These mutations are most commonly located in the NF (V205M, G208R/S, R216Q/W, D218G/N/Y) (11–18), but there are other mutations with similar clinical manifestation located in the region following the CF (insertion of PPFWQ at the position 290, or R307C/H/L, Figure 1A) (19, 20). All of these mutations, which lie within either the NF or close to the CF, may directly influence DNA binding, but no systematic study has yet investigated this aspect on full-length human GATA1 under physiological conditions. In addition to these GATA1 mutations, a splicing defect (intronic mutations or the splice site substitution 332G-C often referred to as V74L), intronic, frameshift mutations, or mutation in the start codon (M1T) lead to the predominant expression of a shorter GATA1 isoform (GATA1s), which lacks the N-terminal transactivation domain (N-TAD), a part of the protein that may also affect the DNA binding properties of GATA1 indirectly (Figure 1A) (21–26). Expression of this short variant, rather than the full-length protein, induces Diamond-Blackfan anemia (DBA) and is implicated in Down syndrome-associated myeloproliferative disorder (transient acute myelopoiesis, DS-TAM), often leading to acute megakaryocytic leukemia (DS-AMKL) (25–27).

**Figure 1.**
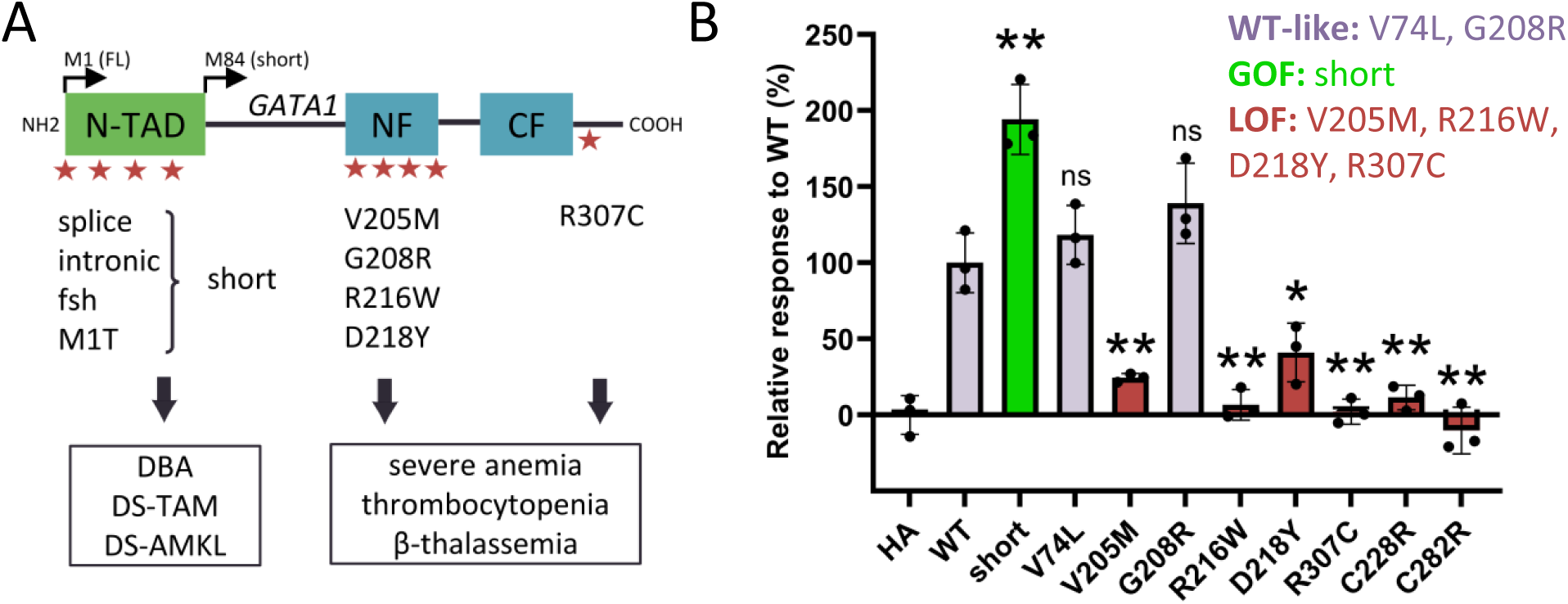
Implication of GATA1 mutations in diseases and functional assessment of their effect on transcription. **A.** The GATA1 mutations characterized in this study. Splicing defects, intronic, frameshift mutations in the first part of the gene or start codon mutation lead to the expression of the short isoform (GATA1s, starting from Met84) which is implicated in Diamond-Blackfan anemia (DBA), Down-syndrome associated transient abnormal myelopoiesis (DS-TAM) or acute megakaryoblastic leukemia (DS-AMKL). Mutations in either in the N-terminal zinc finger (NF) or close to the C-terminal zinc finger (ZF) are implicated in severe anemia, thrombocytopenia and β-thalassemia. **B.** Transcriptional activation function of different mutant forms of GATA1 as measured by luciferase promoter assay using K562 cells and a reporter plasmid containing the full haplotype promoter region of the *ATP2B4* gene. The activity of the different mutant GATA1 proteins in the luciferase assays compared to HA transfected (0%) and HA-GATA1-WT (100%) transfected cells (3 biological and 2 technical replicates, two-tailed t-test on the average of the values measured for biological replicates, ns: non-significant, *: p<0.05, **: p<0.01). Grey: WT-like, green: GOF (gain of function), red: LOF (loss of function). Note, that the expressions were similar for all proteins in K562 cells except for the short isoform, where we detected lower levels of the protein (Supplementary Figure S1). For the raw data see Supplementary Data 1.

Our understanding about the molecular impact of TF mutations on DNA binding is limited due to multiple technical challenges. Most methods used for studying DNA binding are either low-throughput, often requiring extensive recombinant protein production (28, 29), or high-throughput but only qualitative (30, 31). The most commonly used electrophoretic mobility shift assays (EMSA) can provide semi-quantitative insights into binding properties, however, these experiments are labor-intensive, and often rely on non-native conditions, making them unsuitable for the systematic exploration of multiple mutations (32, 33). Recombinant production of full-length TFs can be extremely challenging, as these proteins are difficult to handle, especially when destabilizing pathological mutations are introduced. For this reason, most studies focus only on the DNA-binding domain of GATA1 for biophysical characterization (28, 29). However, this approach can be misleading, as essential regions contributing to binding may be missing. Additionally, for mutations located outside the DNA-binding domain, such as in the case of a short GATA1 variant, this strategy is inadequate. Complementary to these biophysical approaches, ChIP-seq can identify specific binding sites for a given full-length protein genome-wide, however ChIP-seq results are influenced by chromatin structure and it may miss the effects of mutations that cause subtle changes in DNA binding affinity constants (34, 35). Therefore, a versatile technique is needed to characterize the intrinsic properties of DNA-protein interactions to assess how mutations alter these parameters.

Here we adapted our recently developed native holdup (nHU) assay to study the impact of pathological GATA1 mutations on DNA binding (36). Just as the original assay dedicated to protein-protein interactions, this updated version is also suitable for the determination of apparent equilibrium binding constants, which enables the discrimination of stronger and weaker binders, allowing the study of variants that have only a mild effect on DNA binding. A major advantage of this method is that protein purification is not required to quantitatively assess a given DNA-protein interaction, which simplifies the systematic investigation of DNA-binding properties of multiple mutant variants of a TF. As a model for GATA1 recognition sequence, we used the erythroid-specific promoter region of the *ATP2B4* gene, which contains a haplotype that is known to alter the binding of GATA1 (37, 38). We demonstrate that nHU is not only capable of detecting subtle differences in DNA-protein interactions across multiple sequence variants, making it a powerful technique for precisely assessing the binding affinities of GATA1 mutants, but can also be used to identify and simultaneously quantify binding partners of short DNA segments proteome-wide. Using this new version of nHU, we investigated six disease-causing mutations in GATA1 and found that the GATA1s isoform showed markedly increased DNA-binding affinity, while the R307C mutation reduced the affinity of the protein to segments of the *ATP2B4* erythroid promoter. For three other mutations (V205M, R216W, D218Y), although they showed reduced activity in the reporter assay, no change in DNA binding was detected, suggesting that these variants probably affect the regulation through disrupted protein-protein interactions. Integrating data from previous studies, analysis of promoter activation and DNA-binding capacity reveals the distinct functional impacts of different GATA1 mutations.

### Materials and Methods Cell culturing

K562 cells (ATCC ref. CCL-243, RRID: CVCL_0004, cell line was a kind gift of Arnauld Poterszman) were grown in RPMI 1640 medium without HEPES (Gibco) completed with 10% fetal calf serum (FCS) and 40 µg/ml gentamicin and diluted 1:12 every 3rd/4th day. HEK293T cells (authenticated with 100% match as ATCC ref. CRL-3216, RRID: CVCL_0063) were grown in Dulbecco’s modified Eagle’s medium (DMEM, Gibco, glucose: 1 g/liter) completed with 10% FCS and 1% Penicillin-Streptomycin (Gibco) on CellBIND flasks (Corning), diluted 1:10 every 3rd/4th day. All cells were kept at 37°C and 5% CO_2_.

### DNA plasmid constructs and mutagenesis

For the luciferase assay, the pGL3 vector containing the erythroid promoter region of the *ATP2B4* gene with the haplotype was created before (pGL3-ATP2B4-H1st plasmid, haplotype carrying version) (37). For the GATA1 mammalian plasmid, the sequence of GATA1 was cloned from the pIRES2-GATA1-GFP plasmid (37) into the pCI standard mammalian vector containing a HA-tag for N-terminal tagging with standard restriction cloning.

The different mutant forms of GATA1 were created by using standard site-directed mutagenesis on the pCI-HA-GATA1-WT plasmid with Platinum SuperFi II polymerase (Thermo Fisher Scientific). Successful mutagenesis was verified by Sanger-sequencing (Eurofins Genomics Germany GmbH, Ebersberg, Germany).

### Transfections

For the luciferase reporter assay, 1×10^5^ K562 cells/well were seeded the day before the transfection on 24-well plates. The next day, cells were transfected with the pGL3-ATP2B4-H1st-haplo and pCI-HA-GATA1 WT and mutation carrying plasmid vectors using JetPRIME reagent (Polyplus), according to the manufacturer’s recommendations using 0.35 μg pGL3 and 0.15 μg pCI-GATA1 vector DNA and 1 μl JetPRIME reagent/well.

For the cell extract preparation of HA-GATA1 WT and mutants expressing HEK293T cells, the cells were seeded on poly-D-lysine (Sigma-Aldrich, ref. P6407) treated 100 mm cell culture dishes (2×10^6^ cells/dish). The next day, the transfection was carried out with JetPRIME reagent (Polyplus) according to the manufacturer’s recommendations using 5 μg DNA and 20 μl of JetPRIME reagent/well. The medium was replaced with fresh medium after 5 hours of treatment with the transfection mixture.

For the immunostaining of HA-GATA1 expressing cells, 24-well plates with coverslips were poly-D-lysine treated (Sigma-Aldrich, ref. P6407), and HEK293T cells were seeded on the plates the day before the transfection (1×10^5^ cells/well). The transfection was carried out with JetPRIME reagent (Polyplus) according to the manufacturer’s recommendations using 0.5 μg DNA and 1 μl of JetPRIME reagent/well. The medium was replaced with fresh medium after 5 hours of treatment with the transfection mixture to enhance cell survival.

### Luciferase reporter assay

24 hours after transfection, K562 cells were collected by centrifugation (1 min x 1000g) and washed once with PBS. Then, 100 μl of 1x passive lysis buffer (5x PLB diluted with UltraPure water, Promega ref. E1941) were added to the cells and incubated for 15 minutes at room temperature under shaking. Lysates were pipetted up and down a few times and 20 μl lysate was transferred into 96-well white plates (Millipore ref. MSSWNFX40). Then, 100 μl luciferase assay reagent (Promega, ref. E1483) was added to each well and the firefly luminescence was measured immediately on a PHERAstar (BMG Labtech, Offenburg, Germany) microplate reader by using Lum Plus filter. The remaining cell lysates were kept for expression determination by Western blot. In total, 3 biological (transfection) and 2 technical (from the same extract) replicates were performed. For statistics, the technical replicates were averaged and two-tailed t-tests were performed on these averaged values for each biological replicate (n=3). For statistical analyses, GraphPad Prism 10 software was used. For raw data and calculations see Supplementary Data 1.

### Cell extract preparation for nHU assay

Cell extracts were prepared similarly as before (36) except the lysis buffer composition, which was optimized for DNA-protein interactions: 50 mM Hepes-KOH (pH 7.5), 100 mM NaCl, 50 mM KCl, 4 mM MgCl_2_, 1% Triton X-100, 5× cOmplete EDTA-free protease inhibitor cocktail, 5 mM TCEP, and 10% glycerol. Briefly, K562 cells were collected by centrifugation (1000g × 5 min), quickly washed with PBS, and ice-cold lysis buffer was added. HEK293T cells were washed once with PBS, lysis buffer was added and cells were collected by scraping. Lysates were sonicated (4×20 s with 1-sec-long pulses on ice) and incubated rotating (4°C for 30 min). The lysates were centrifuged (12,000 g 4°C for 20 min), and the supernatant was kept. Total protein concentration was measured by standard Bradford method (Bio-Rad protein assay dye reagent, ref. 5000006) using a bovine serum albumin (BSA) calibration curve (MP Biomedicals, ref. 160069) on a UV- 1700 PharmaSpec spectrophotometer instrument. Extracts were diluted to 1 mg/ml concentration and were snap-frozen in liquid nitrogen and stored at −80°C.

### Biotinylated DNA oligonucleotides and annealing for nHU

All DNA oligonucleotides were produced by Eurofins Genomics (Eurofins Genomics Germany GmbH, Ebersberg, Germany) and were high-performance liquid chromatography (HPLC) purified. The following oligonucleotides were used (5’-3’ orientation, GATA1 recognition sequences and their haplotype versions are labelled green, Biot: biotin-group):

4945_WT:

For: Biot-TCAGCCCTCCG**TATC**GTCACCTACAC

Rev: GTGTAGGTGAC**GATA**CGGAGGGCTGA

4945_HAP:

For: Biot-TCAGCCCTCCG**TTTT**GTCACCTACAC

Rev: GTGTAGGTGAC**AAAA**CGGAGGGCTGA

51_WT:

For: Biot-GAATTGAGAGG**TATC**T**TATC**GCTCCCACTCC Rev: GGAGTGGGAGC**GATA**A**GATA**CCTCTCAATTC

51_HAP:

For: Biot-GAATTGAGAGG**TATC**T**TACC**GCTCCCACTCC

Rev: GGAGTGGGAGC**GGTA**A**GATA**CCTCTCAATTC

SCR1:

For: Biot-CCACTCTGTTCTAGAGAAGCCTTCGTGAACT

Rev: AGTTCACGAAGGCTTCTCTAGAACAGAGTGG

SCR2:

For: Biot-ATGCCATTGCTTCATGCACGTATAGCCACTG

Rev: CAGTGGCTATACGTGCATGAAGCAATGGCAT

CTRL_palindrome_:

For: Biot-GCGC**TATC**A**GATA**AGGCCTTG

Rev: CAAGGCCT**TATC**T**GATA**GCGC

For annealing, the two single-stranded oligonucleotides were mixed in 1:1.5 forward-reverse DNA ratio in a 12.5 μM final double-stranded oligonucleotide concentration in holdup (HU) buffer (50 mM Tris pH 7.5, 300 mM NaCl, 1 mM TCEP, 0.22-μm filtered). In a pre-heated thermoblock the samples were heated up to 95 °C for 5 minutes, then slowly cooled down within 100 minutes until they reached room temperature.

In general, 680 μl of 12.5 μM dsDNA were used for the saturation of 25 μl pure streptavidin resin (Streptavidin Sepharose High Performance, Cytiva). The estimated bait concentration of the saturated resin is 10 μM once 25 μl pure bait-saturated streptavidin resin was mixed with 100 μl of cell extract.

### Singe point and titration native holdup

For saturating resin (Streptavidin Sepharose High Performance, Cytiva) with dsDNA or biotin (as control), 25 μl pure resin was mixed with either 680 μl of 12.5 μM dsDNA or 25 μl 1 mM biotin, and additional NaCl was added (at 0.5 M final concentration) to facilitate DNA fixation to the resin. Resin mixtures were incubated for 1 hour at room temperature, gently mixing every 15 minutes. Resins were briefly centrifuged (15 s, 2000 g), the supernatants were removed and the resins were depleted with biotin (10 resin volume, 3 min, holdup buffer supplemented with 100 μM biotin). Then, resins were washed four-times (10 resin volume, holdup buffer). To get a bait concentration of 5 μM, these pre-saturated resins were reconstituted in 6x resin volume of HU buffer and were mixed in a 1:1 ratio by adding 12.5 μl dsDNA to 12.5 μl control (biotin) pure resin. Finally, all liquid was removed and cell extracts were added in 1 mg/ml concentration in 1:4 pure resin:extract ratio (25 μl pure resin: 100 μl cell extract) and incubated for 2 hours at 4 °C with constant gentle agitation. After the incubation ended, the resins were separated from the supernatant by a brief centrifugation (15 s, 2000 g). 75% of the supernatants were quickly pipetted to a separate tube and centrifuged again briefly, and 50% of the original volume of the supernatant was kept in the end to reduce resin contamination in the final samples.

Titration experiments (coupled to Western blot) were carried out similarly as before, by mixing control and bait-saturated resins and keeping the total resin volume constant (36). Twelve-point titrations were carried out. For the starting point, the DNA-saturated and biotin control resins were mixed in 1:1 (5 μM starting point) or 1:3 (2.5 μM staring point) ratio, typically 20+20 μl or 10+30 μl, respectively, calculated for the pure resin volume, in 4x resin volume in holdup buffer. Then, 1:1 serial dilution was carried out with the control resin (20+20 μl pure resin). The final tube contained only the biotin saturated resin as control. The resins were then briefly centrifuged (15 s, 2000 g), the supernatant was removed and cell lysates were added at a concentration of 1 mg/ml or 0.1 mg/ml, and further procedures were carried out as described for the single-point experiments (20 μl of resin was mixed with 80 μl of extract to obtain the correct concentration of bait during the incubation). All the experiments were carried out in Protein LoBind tubes (Eppendorf, refs. 022431064 and 022431081). All titrations were carried out with minimum two replicates.

### Sample digestion and mass spectrometry (LC-MS/MS) analysis

MS analyses were performed as described before (36, 39). Briefly, nHU samples were precipitated with TCA, and the urea-solubilized, reduced and alkylated proteins were digested with trypsin at 2 M final urea concentration. Peptide mixtures were then desalted on C18 spin-column and dried on Speed-Vacuum. 100 ng peptide mixtures were analyzed using an Ultimate 3000 nano-RSLC coupled in line, via a nano-electrospray ionization source, with an Orbitrap Exploris 480 mass-spectrometer (Thermo Fisher Scientific, Bremen, Germany) equipped with a FAIMS (high Field Asymmetric Ion Mobility Spectrometry) module. Data was collected in DDA (data dependent acquisition) mode, proteins were identified by database searching using SequestHT (Thermo Fisher Scientific) with Proteome Discoverer software (Thermo Fisher Scientific). Peptides and proteins were filtered with a false discovery rate (FDR) at 1%. Label-free quantification was based on the extracted ion chromatogram (XIC) intensity of the peptides. All samples were measured in three technical triplicates. The measured XIC intensities were normalized based on median intensities of the entire dataset to correct minor loading differences. For statistical tests and enrichment calculations, not detectable intensity values were treated with an imputation method, where the missing values were replaced by random values similar to the 10% of the lowest intensity values present in the entire dataset. Unpaired two tailed t-tests, assuming equal variance, were performed on obtained log_2_ XIC intensities. All raw LC-MS/MS data have been deposited to the ProteomeXchange via the PRIDE database with identifier PXD055181. Obtained fold-change values were converted to apparent affinities using the hyperbolic binding equation and binding thresholds at 2σ were determined as described before in details (36).

### Gene Ontology enrichment analysis

Gene Ontology enrichment analysis was performed using the DAVID tool with default parameters (40). Two independent analysis were performed using all the preys that were interacting with at least one bait, or using only those preys that were interacting with all baits, and both analyses were done against the same background (all proteins detected by the proteomic measurements). Only the Cell Component (CC_DIRECT) and Molecular Function (MF_DIRECT) terms were analyzed and all significant terms are included in Supplementary Data 3.

### Identification of sequence-specific binders based on cumulative Δp*K*_app_

To calculate affinity differences between interactions measured with the WT and haplotype-carrying dsDNA baits, we used the apparent affinities, that were converted from the measured fold-change values. First, we identified all transcription factors among the interaction partners of the dsDNA baits using a recently reported comprehensive transcription factor catalog (41). Then, we collected the apparent affinities of all these proteins and calculated the overall effect of the haplotype by summing the affinity differences between WT and HAP sequences. For getting the affinity differences, we simply subtracted the measured p*K*_app_ value of the haplotype sequence from the measured p*K*_app_ value of the WT sequence, that is proportional to the ΔΔ*G*_app_ values (39). For the affinity matrix for WT and haplotype sequences see Supplementary Data 3.

### Western blot

The samples were mixed with 4× Laemmli buffer (120 mM Tris-HCl pH 7, 8% SDS, 100 mM dithiothreitol, 32% glycerol, 0.004% bromphenol blue, and 1% β-mercaptoethanol) in a 3:1 ratio. Equal amounts of samples (generally 6 μg) were loaded on 12% acrylamide gels. Transfer was done using a Trans-Blot Turbo Transfer System and Trans-Blot Turbo RTA Midi 0.45 µm LF PVDF Transfer Kit (Bio-Rad, ref. 1704275). After 1 hour of blocking in 5% milk in TBS-Tween (1x Tris-buffered saline, 0.1% Tween 20), membranes were incubated overnight at 4°C in primary antibody in 5% milk in TBS-Tween. The following antibodies and dilutions were used: anti-GATA1 (1:10,000, Abcam, ref. ab181544, RRID: AB_2920794), anti–glyceraldehyde-3-phosphate dehydrogenase (GAPDH, 1:5,000, Sigma-Aldrich, ref. MAB374, RRID: AB_2107445), and anti-HA.11 epitope tag (1:5,000, BioLegend, ref. 901501, RRID: AB_2565006). Membranes were washed three times with TBS–Tween and incubated at room temperature for 1 hour in secondary antibody solution [Jackson ImmunoResearch, peroxidase-conjugated Affinipure goat anti-mouse (H + L), ref. 115-035-146 RRID: AB_2307392 and goat anti-rabbit (H + L), ref. 111-035-003 RRID: AB_2313567] in 5% milk (concentration 1:10,000). After washing three times with TBS-Tween, membranes were exposed to chemiluminescent horseradish peroxidase substrate (Immobilon, ref. WBKLS0100) and photographed in a docking system (Amersham ImageQuant 800). Densitometry was carried out on raw Tif images by using Fiji ImageJ 1.54f. Between different primary antibody labeling, the membranes were either exposed to 15% H_2_O_2_ to remove secondary signal (in the case of different species) or stripped with mild stripping buffer (15 g/liter glycine, 1 g/liter SDS, and 1% Tween 20, pH 2.2) to remove primary signal (in the case of same species). For native holdups, GAPDH was used as a non-binding control, but was not used to normalize GATA1 values as this induces technical errors coming from blotting issues rather than from differences between the samples, as discussed before (36). In case of inconsistencies in GAPDH values, the blot was repeated.

### Immunostaining of HA-GATA1 transfected cells and colocalization analysis

24 hours after transfection, HEK293T cells were washed once with PBS and fixed with 4% PFA (Electron Microscopy Sciences, ref. 15710) in PBS for 10 minutes at room temperature. The cells were then washed three times with PBS and permeabilized with 500 μl/well of 0.2% Triton X-100 (Sigma-Aldrich, ref. T8787) in PBS for 10 minutes at room temperature. After permeabilization, the samples were blocked in blocking buffer [0.05 g/mL BSA (MP Biomedicals, ref. 160069), 0.1% Triton X-100 in PBS] for 1 hour at room temperature. Then, the cells were incubated overnight with the HA primary antibody (anti-HA.11 epitope tag, dilution: 1:1,000, BioLegend, ref. 901501, RRID: AB_2565006) at 4 °C in blocking buffer. The next day, the cells were washed three-times with PBS with 10 minutes incubations each time at room temperature. Then, the cells were incubated with the secondary antibody (Alexa Fluor 594–conjugated anti-mouse, dilution: 1:1000; Invitrogen, ref. A-11032, RRID: AB_2534091) in blocking buffer. After three-times 10 minutes PBS washes, the coverslips were mounted to microscopy slides by Vectashield mounting medium with 4′,6-diamidino-2-phenylindole (DAPI) (Vector laboratories, Burlingame, CA). Images were taken using a Leica SP5 confocal microscope (Leica Camera AG, Wetzlar, Germany) with an HCX PL APO 63×/1.40 to 0.60 oil objective using excitation at 405 nm (diode) and 594 nm (HeNe laser) and emission at 415 to 480 and 610 to 695 nm for DAPI and Alexa 594, respectively. Images were processed by the Fiji ImageJ 1.54f software. To determine colocalization of HA-GATA1WT and mutant proteins, Coloc2 plugin was used. For each image, the background was subtracted, and three ROIs of the same size were selected, excluding non-transfected cells, large background areas, and dividing cells. After ROI selection, the Coloc2 plugin was used with the Costes automated threshold to determine Manders’ tM1 (MCC), representing the fraction of HA signal in compartments containing DAPI. The three values obtained for each image were then averaged. In total, four images per GATA1 variant were statistically probed using two-tailed Student’s t-test (n=4).

## Results

### Characterization of the functional effects of GATA1 mutations on the *ATP2B4* alternative promoter using luciferase assay

In order to shed light on the functional effects of different disease-associated GATA1 mutations on transcription, we performed a luciferase reporter assay in K562 cells. As a reporter construct, we used the alternative erythroid specific promoter of the *ATP2B4* gene followed by the Firefly luciferase. This erythroid promoter is located just upstream of the second exon of the gene, and drives the expression of the PMCA4b calcium pump specifically in erythroid lineages (38, 42). In this erythroid promoter, a haplotype has been identified that causes lower expression of the protein in mature red blood cells, reduces mean corpuscular hemoglobin concentration in the blood and provides certain degree of protection against malaria (38, 43–46). We showed in our previous study that a significant increase in transcription can be detected once GATA1 is co-transfected along with the reporter plasmid containing this erythroid promoter and that the largest window in the signal difference could be obtained with the haplotype version of the promoter, that still contains eight predicted single GATA1 binding sites (37). We used this haplotype-containing reporter plasmid along with a HA-GATA1 construct, containing either the full-length WT or the mutant GATA1 sequences in the luciferase assay (Figure 1, Supplementary Data 1). A total of 10 different GATA1 constructs were generated, including the WT sequence, the short variant (GATA1s, or simply short) lacking the N-TAD, the V74L, which serves as a control since it does not exist in its natural form, as the short isoform is expressed as a result of this mutation, NF mutations (V205M, G208R, R216W, D218Y), the R307C mutation that is located just downstream the CF, and finally two artificial mutations disrupting the DNA binding of either the NF (C228R) or the CF (C282R) by eliminating one of the cysteines coordinating the zinc ion. After 48 hours of transfection we lysed the cells and measured the amount of luciferase produced based on luminescence. The remaining lysates were subjected to Western blot analysis to determine the GATA1 expression levels for each condition (Supplementary Figure S1, Supplementary Data 1).

Expression levels of all GATA1 mutants were similar to WT GATA1, except for the short variant, which expressed at a significantly lower level (Supplementary Data 1, Supplementary Figure S1). Most artificial and pathological mutations caused loss of function, based on the luciferase activity. For the artificial C228R and C282R mutations, which disrupt the folding of the NF and CF DNA-binding domains, respectively, we observed a complete loss of function in the luciferase assay, emphasizing the critical role of both zinc fingers in maintaining function (Figure 1B). Similarly, the R216W and R307C mutants showed complete loss of function. Partial loss of function was observed for the V205M and D218Y mutations, which are located within the NF region of the protein. No functional impairments were detected with the artificial V74L and the natural G208R mutants, both of which exhibited luciferase activity comparable to the WT. Interestingly, the short GATA1 variant showed an almost twofold increase in activity compared to WT GATA1, even though Western blot analysis indicated that cells expressed about half the amount of this variant compared to WT, suggesting an approximately fourfold increase in partial activity for this mutant when normalized to GATA1 expression (Figure 1B, Supplementary Figure S1).

### Proteome-wide affinity characterization of the *ATP2B4* promoter elements by native holdup

The luciferase promoter assay revealed a set of GATA1 mutations that cause perturbed activity, yet it does not shed light on the mechanism behind these changes. Further investigations are necessary to decipher if these mutations affect primarily the DNA binding or act through other functions of the protein, such as essential protein-protein interactions (10, 15). To investigate further the molecular mechanism without having to produce large quantities of all studied GATA1 variants for biophysical assays, we decided to adapt the native holdup (nHU) method to DNA-protein interactions to investigate the effects of different mutations on DNA binding. This technique is advantageous because it is simple, fast, provides quantitative affinity measurements, and does not require protein purification. This is a comparative chromatographic approach where a cell extract (analyte), is mixed with a bait-or control-saturated resin and the unbound prey protein fraction is measured from the liquid phase after incubation and rapid separation of the liquid and solid phases (Figure 2) (36, 47–49). From this unbound fraction value and from the estimated bait concentration during the experiment, the affinity of the bait-prey interaction can be calculated with high precision and accuracy.

**Figure 2.**
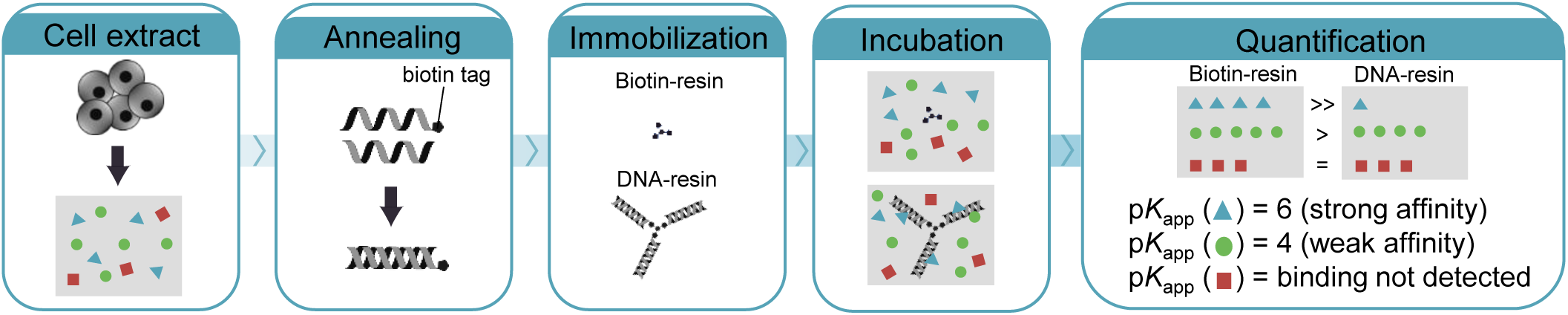
The principles of native holdup using biotinylated oligonucleotide baits. The first step is to produce total cell extracts. Then, annealing is performed using one biotinylated and one non-biotinylated ssDNA molecule. The annealed dsDNA is immobilized on streptavidin resin along with a control (biotin-treated) resin. These saturated resins are then incubated with the cell extract until the equilibrium is reached, after which the supernatant is rapidly separated and quantification is performed from the liquid phase. Finally, the unbound fractions are calculated by comparing the reaction containing dsDNA to the control (biotin-containing) reaction and converted to apparent affinity using the hyperbolic binding equation.

In order to validate the applicability of the native holdup assay for DNA-protein interactions, we investigated the affinity interactomes of the different *ATP2B4* erythroid-specific alternative promoter elements by native holdup coupled to mass spectrometry (nHU-MS, Figure 2). In our previous studies, we performed the assay at relatively high bait concentrations to capture weak and transient protein-protein interactions (36). However, here we had to consider the nature of nucleic acid-protein interactions, which typically have a higher affinity than protein-protein interactions, and these interactions greatly depend on ionic strength. For these reasons, we performed the nHU experiments at a lower estimated bait concentration (5 µM), using a more dilute cell extract (1 mg/ml) that the one we used for protein-protein interaction assays. We have also optimized cell lysis conditions to make it more optimal for capturing nucleic acid-protein interactions by the addition of Mg^2+^ and K^+^ ions and by slightly lowering the ionic strength of the lysis buffer.

We have previously identified two SNP-containing sites in the alternative promoter region of the *ATP2B4* gene, referred to here as 4945 and 51 (abbreviations for SNP identification codes rs10751449/rs10736845 and rs10751451), which are single and tandem GATA1 recognition sites altered by the haplotype (38). In reporter assays where this erythroid promoter was inserted to drive the expression of a luciferase, the mutation of any or both of these SNPs was enough to recapitulate the reduced expression observed with the full haplotype sequence (37). We used the wild-type and haplotype-containing double-stranded (ds) DNA constructs for the 4945 and the 51 fragments (26-mer and 31-mer, respectively) that we hybridized before the nHU experiment (Figure 3A). Out of the synthetic single-stranded (ss) DNA constructs, only forward strands were biotinylated. To capture the interactions present in erythroid cells, K562 cell extracts were prepared. Although we considered using a nuclear extract, we hypothesized that the depth of standard MS measurements is going to be sufficient to capture most interactions of abundant nuclear proteins from total cell extracts and this way we do not introduce any sort of technical bias in our experiment. Apart from these minor modifications, we followed the original nHU protocol with biotinylated-dsDNA fragments as baits against biotin controls (Figure 2). After incubating the bait-saturated resins with cell extracts until equilibrium was reached, we collected the supernatant and analyzed it using routine label-free quantitative mass spectrometry. In these total proteomic measurements, we managed to quantify the relative amounts of 4,560 proteins (Figure 3B, Supplementary Data 2). Binding threshold was set as discussed in details before. Briefly, a hyperbolic threshold was used where significance had to reach a probability of at least 0.05, calculated by a two-tailed unpaired t-test, and prey concentration change needed to be greater than 95% of the measured changes (>2 σ) (36). Based on this threshold, 274 proteins showed significant binding to at least a single bait (Supplementary Data 2).

**Figure 3.**
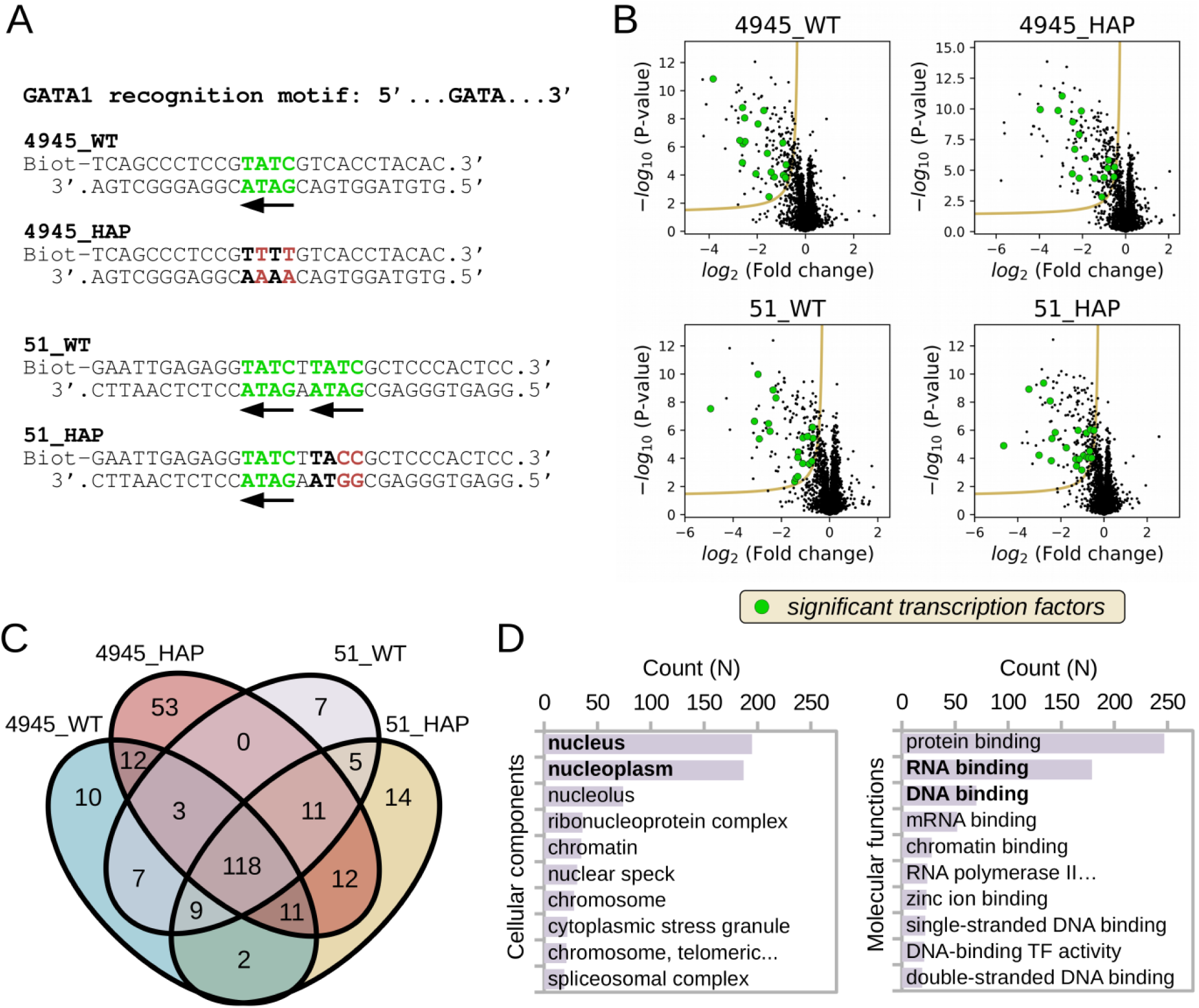
Affinity interactome determination of *ATP2B4* erythroid-specific alternative promoter elements measured by mass spectrometry-coupled nHU on K562 cell extracts. **A.** Oligonucleotides used in the native holdup assay. Both WT and haplotype (HAP) sequences were used for regions 4945 and 51. For 4945, the single GATA sequence is abolished by the haplotype, whereas for the tandem region 51, one GATA site is eliminated by the haplotype. **B.** Volcano plots of four experiments with the four different baits (n=3, injection replicates). Transcription factors are highlighted in green. For further data see Supplementary Data 2. **C.** Overlap of the interaction partners of the different baits, visualized with a Venn diagram (hyperbolic significance threshold was determined at 2σ). Surprisingly, most of the partners found were detectable with all four baits. **D.** Gene ontology enrichment was performed using the DAVID tool (40) on the proteins that were binding to at least one DNA bait. Most of the partners detected are general nucleic acid binding proteins that are located in the nucleus. For further data see Supplementary Data 3.

### GATA1 is a sequence-specific partner of the *ATP2B4* promoter elements measured by native holdup

Substantially more partners were detected with the DNA baits than in the case of performing native holdups with peptide baits, which is surprising given the short size of the nucleic acids used in the assay (36, 50). Similar numbers of partners were only detected when promiscuous globular domains (e.g. SH3 or PDZ) or full-length proteins were used as baits (36, 39). A possible explanation for the large number of interaction partners is that nHU can capture indirect interactions. When large molecular machines are captured, all their subunits are identified as interaction partners (36). However, of the interactions identified, only a few large complexes have been identified, such as the RNA exosome complex, the RNA polymerase III complex, or components of the spliceosome, covering only a small fraction of all detected partners (Supplementary Data 2). Additionally, the different dsDNA-bait fragments showed remarkably similar interactome in qualitative terms. Among the 274 partners, 118 (43%) were identified as significant partners with all four dsDNA baits, while 152 (55%) and 190 (69%) proteins were identified as partners with three and two baits, respectively (Figure 3C).

In order to interpret the results of the nHU-MS experiments, we used the DAVID tool to perform Gene Ontology enrichment analysis for cellular compartments and molecular functions (Figure 3D, Supplementary Data 3) (40). Considering all 274 partners, we have found a great enrichment of proteins localized in the nucleus (N=190; P = 2.0 x10^-44^) or more precisely in the nucleoplasm (N=183; P = 1.4 x10^-67^). These partners were also greatly enriched with nucleic acid binding proteins. However, surprisingly, we identified more RNA binding proteins (N=177; P = 5.0 x10^-129^) than DNA binding proteins (N=73; P = 7.4 x10^-24^). Looking more closely at the 118 proteins that we identified as partners with all dsDNA baits, we also found that nuclear proteins (n=89; P = 5.4 x10^-25^), RNA binding proteins (N=79; P = 1.1 x10^-58^), and DNA binding proteins (N=37; P = 9.1 x10^-15^) were highly enriched. Not surprisingly, we also identified many general DNA-binding proteins among the partners, such as multiple linker histone variants. Therefore, we concluded that the affinity interactomes of nucleic acid baits can be characterized exhaustively by simultaneously measuring sequence-specific and sequence-independent interactions using nHU.

Sequence-independent interactions are general properties of nucleic acids, related to their net charge and conserved topology. In sequence-independent interactions, molecular contacts are mostly made with the phosphate backbone and the contribution of the nucleic acid sequence is marginal. Focusing only on TFs (41), we identified 32 TFs among the 274 proteins showing binding to at least one of the dsDNA baits (accounting for 11.7%, Supplementary Data 1). This is apparently a slightly higher ratio than the general occurrence of TFs in the entire genome (1,639 TFs among the 20,428 human proteins, accounting for 8.0%), and a much higher ratio than the measured occurrence of TFs in the explored portion of the erythroid proteome of K562 cells (141 detected TFs out of 4,560 detected proteins, accounting for 3.1%). However, even TFs can bind indirectly, or in a sequence-independent manner, similarly to other nucleic acid binding proteins discussed above. To identify the TFs that bind in a sequence-specific way, we exploited the quantitative aspect of the nHU assay by comparing the measured apparent affinities of the wild-type and haplotype dsDNA baits. By using an unbiased approach, we expected either a gain of affinity or a loss of affinity upon the SNPs. Additionally, since both sites similarly contribute to the regulation of *ATP2B4* gene expression, we assumed that the same interaction partners would have similar effects on both sites. Only a few TFs showed increased affinity for both haplotype baits, such as LIN28B or ATF2, but these showed minor changes in affinity rather than large perturbations (Table 1). Multiple TFs showed decreased affinities to the 51 site, such as ZNF24, or HOXB4, but these proteins did not show detectable binding to the 4945 site. Considering the 4945 segment, the interaction partner showing the highest down-regulation in affinity was GATA1. GATA1 also showed binding to the 51 site that displayed decreased affinity to the haplotype sequence, albeit only a slight change was measured. Nevertheless, upon ranking of all transcription factors based on their cumulative Δp*K*_app_ values of these sites (which is related to ΔΔG), GATA1 showed the highest overall affinity change out of all partners (Table 1). From these observations, we concluded that GATA1 is indeed the sequence-specific regulator in this region, consistent with previous findings.

**Table 1.**
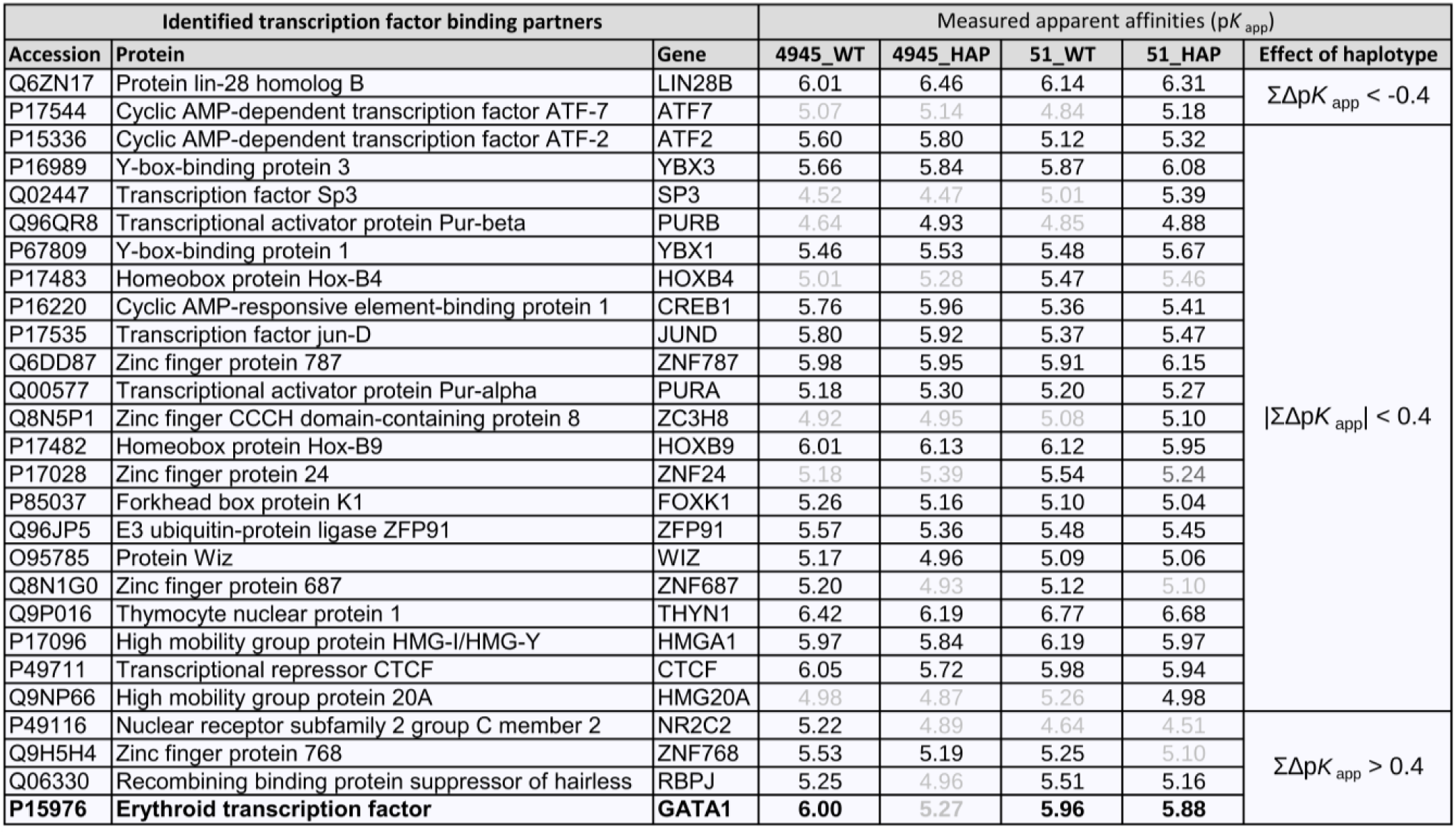
Identification of sequence-specific binders by calculating cumulative Δp*K*_app_ for transcription factors detected in the mass spectrometry coupled nHU experiments. Apparent affinities that belong to interactions below the significance threshold are colored in grey. To calculate the effect of the haplotype, the WT and haplotype (HAP) apparent affinity differences were summed and the measured p*K*_app_ value of the haplotype sequence was subtracted from the measured p*K*_app_ value of the WT sequence. GATA1 is highlighted in bold as it showed the largest negative effect due to the haplotype.

### *Ab ovo* identification of sequence-specific interactions with nHU

By using an unbiased approach, we could demonstrate that the nHU assay is well-suited for identifying sequence-specific interactions. Exploiting clinically relevant variants allowed us to distinguish relevant interactions from sequence-independent or irrelevant interactions. Yet, this approach could not clearly separate GATA1 from other transcription factors, leaving the possibility that other interaction partners could also contribute to the differential *ATP2B4* gene expression. Moreover, this approach would have failed to identify GATA1 as the critical partner of the 51 site alone, that only showed small apparent affinity change in the single point affinity measurement. To overcome these issues, we decided to use scrambled dsDNA baits, instead of free biotin, to saturate the control resin stock. By comparing binders against this control scrambled dsDNA bait, we can identify all sequence-specific interaction partners of a given oligonucleotide bait, irrespective of mutations or polymorphisms within the DNA fragment.

Thus, we repeated the nHU-MS experiments with the 51 WT fragment but using two additional scrambled DNA fragments as controls (Figure 4A-D, Supplementary Data 4). A major limitation that we faced during this assay, that due to detection limitations of mass spectrometry, we failed to detect GATA1 in the K562 proteome. For this reason, we performed Western blot experiments using the same samples that we also measured with proteomics and with the same experimental strategy (same number of replicates, same statistical analysis) to quantify GATA1 protein specifically (Figure 4E, Supplementary Figure S2). Again, we detected a high number of interactors for all the three DNA probes compared to the biotin-saturated resin controls, but only in the case of 51 WT sequence we detected GATA1 as a binding partner (Figure 4B-D). In total, we had 79 partners that were found to interact with all three dsDNA baits and 43 further proteins found to bind two of the dsDNA baits (Figure 4F). Therefore, just as in the first experiment, the measured interactomes of the different dsDNA baits were similar and we only found 16 specific binding partners for the 51 sequence.

**Figure 4.**
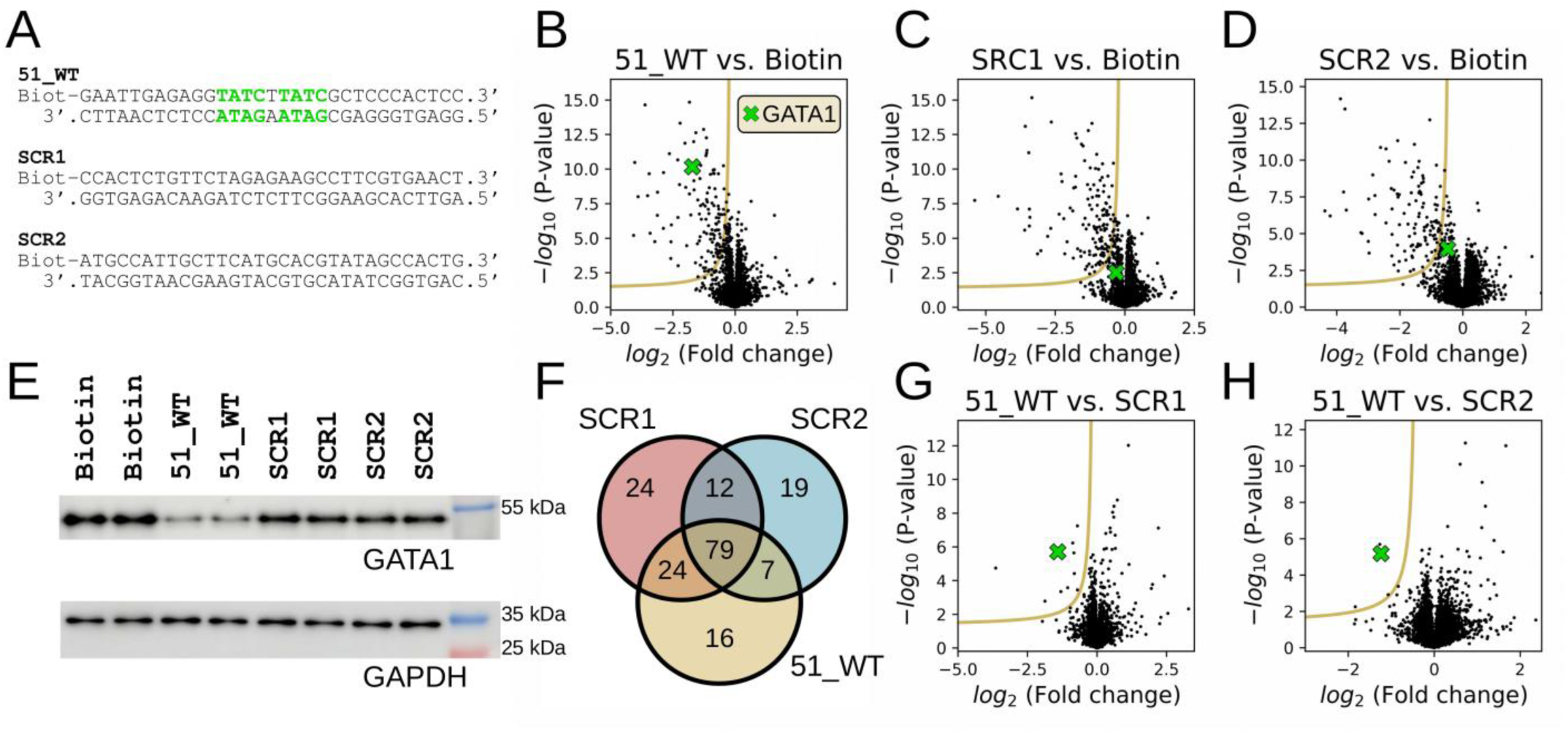
*Ab ovo* identification of sequence-specific interactions for a given DNA-oligonucleotide with nHU coupled to mass spectrometry. **A.** Oligonucleotides used in the experiment series. We used the 51_WT sequence and two scrambled versions (SCR1 and SCR2) to find the sequence specific recognition partners for the 51_WT bait in an unbiased manner. The GATA sequences are highlighted in green. **B.** Volcano plot visualization of the nHU result with 51_WT sequence compared to the biotin control. **C.** Volcano plot visualization of the nHU result with SCR1 sequence compared to the biotin control. **D.** Volcano plot visualization of the nHU result with SCR2 sequence compared to the biotin control. All native holdup experiments were carried out with 2 biological replicates and their analytics were done with 3 technical replicates for each biological replicate (n = 2×3). GATA1 values were determined by Western blots from the same samples and highlighted in green in each volcano plots. **E.** Western blot analyses of GATA1 levels in the native holdup reactions subjected to mass spectrometry analyses in B-D (2 biological replicates and 3 technical replicates for each biological replicate, n = 2×3, for full blots see Supplementary Figure S2). GAPDH was used as non-binding loading control. **F.** The number of detected significant partners for each oligonucleotide baits visualized with a Venn diagram. Significance was determined based on hyperbolic significance threshold at 2σ. **G.** Volcano plot visualization of the native holdup results with the 51_WT bait compared to SCR1 bait. **H.** Volcano plot visualization of the native holdup results with the 51_WT bait compared to SCR2 bait. In both cases, GATA1 (green) has been identified as a significant sequence-specific binder (at 2σ). For further data see Supplementary Data 4.

When we used the SCR1 or SCR2 separately as control samples, we only detected 10 or 4 statistically significant binding partners, respectively, for the 51 WT sequence (Figure 4G-H, Supplementary Data 4). Among them, only 3 were common in both comparisons, namely TMEM258, CNBP and GATA1. TMEM258 is a transmembrane protein that is a common contaminant in mass spectrometry measurements, so it is likely that this protein was detected as a partner due to its nonspecific binding to our probe. CNBP is a CCHC-type seven zinc finger nucleic acid binding protein that preferentially binds single-stranded oligonucleotides. The predicted binding motif consensus is somewhat consistent with our 51 WT sequence (51, 52). Due to this, we believe that our annealing was not completely efficient, or considering that the collective electrostatic binding of seven zinc fingers may increase the overall affinity to DNA due to avidity, the protein may appear as a sequence-specific binder in this study. Finally, GATA1 was the only protein that is a transcription factor and also binds to dsDNA. Thus, we were able to identify the relevant transcription factor that binds to our target sequence even without the use of sequence-disrupting mutations. Importantly, GATA1 shows minimal binding to scrambled DNA sequences, indicating that it does not bind strongly to DNA baits in a sequence-independent manner under native conditions. Therefore, most of the observed interactions between the various *ATP2B4* baits and GATA1 detected in the nHU-MS experiment are driven by sequence-specific interactions.

### Biochemical characterization of the interactions between the *ATP2B4* promoter elements and GATA1

Although the single point nHU experiment used in combination with mass spectrometry provides a proteome-wide estimation of the affinity interactome of a given oligonucleotide, it lacks certain biochemical features. Most importantly, to convert the measured fractional prey depletion to apparent affinity, we had to assume a simple binding mechanism with no complications. In order to determine precisely the affinities, we performed titration nHU experiments and monitored GATA1 depletion from the liquid phase with Western blot (Figure 5A, Supplementary Figure S3). We found that both WT *ATP2B4* fragments displayed simple binding mechanism (matching a hyperbolic binding equation) but they also showed a partial GATA1 binding activity of about 75%, meaning that 25% of the endogenous GATA1 is unable to interact with the bait DNA fragments even at high bait concentrations (Figure 5B). We also found that the fragment 51, that contains two consecutive GATA motifs in tandem orientation, mediates higher affinity interactions with GATA1 with a dissociation constant of 17 nM, while the fragment 4945, that contains a single GATA motif, mediates a somewhat weaker affinity interaction with GATA1 with a dissociation constant of 100 nM (Figure 5B). By repeating the titration experiment with the haplotype sequences, we found that both promoter fragments displayed a weaker interaction. The haplotypic version of fragment 51, which still contains a single GATA motif, showed a dissociation constant of 130 nM with GATA1, which is about 10-fold weaker than that of the WT 51 sequence, and similar to the affinity measured for WT 4945, which also contains a single motif. The haplotype version of fragment 4945, that do not contain any GATA motif, displayed only a marginal interaction, approximately 100-times weaker than the WT sequence (Figure 5B), possibly due to marginal sequence-independent binding. Thus, by performing native holdup titration experiments, we can precisely determine the equilibrium binding constants of DNA-protein interactions and detect even small differences in affinities, which is essential for characterizing the effects of transcription factor mutations.

**Figure 5.**
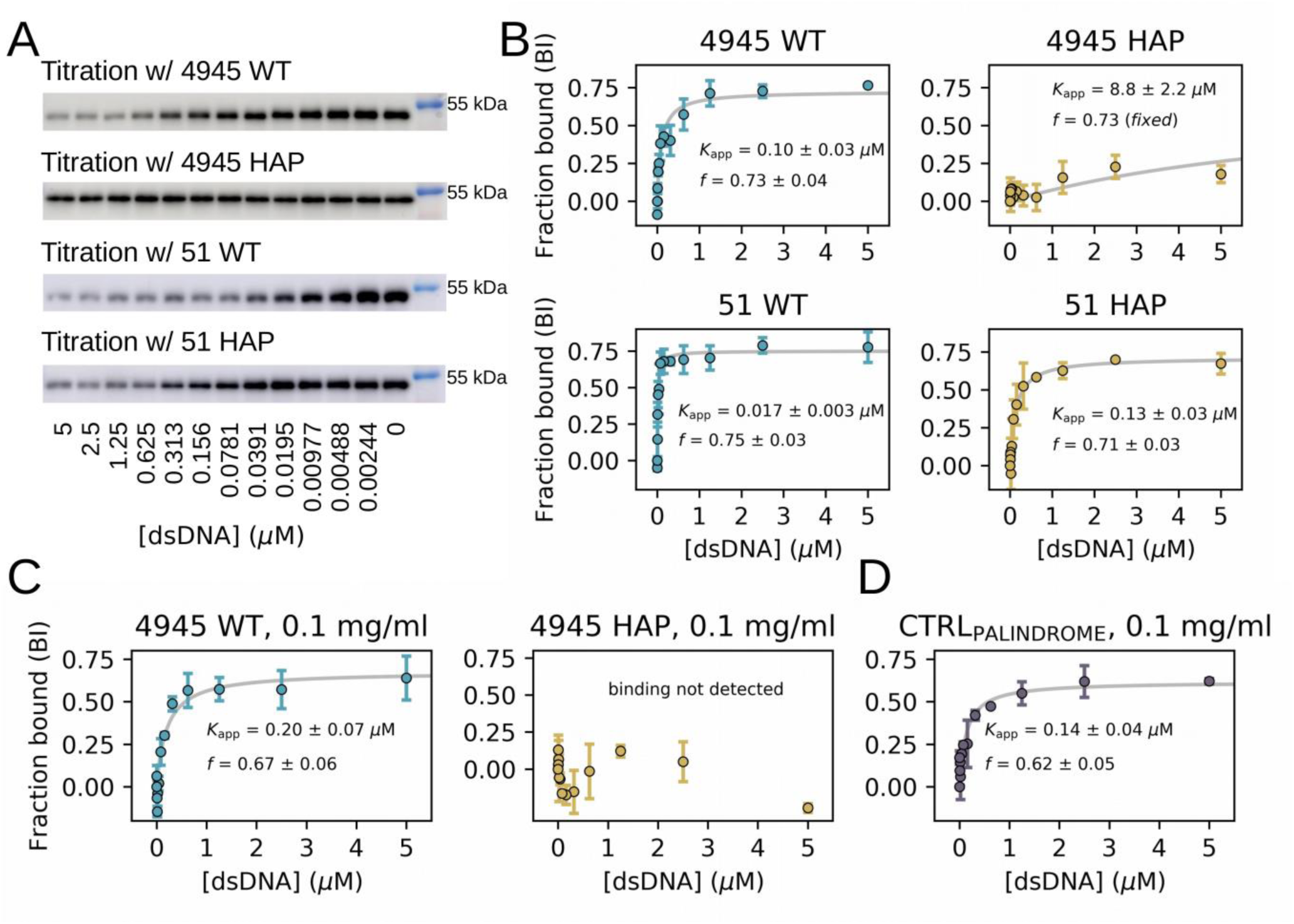
Biochemical characterization of DNA binding of GATA1 to the WT and haplotype-carrying *ATP2B4* promoter elements using nHU titration coupled to Western blot on K562 cell extracts. **A.** Western blot results of the titrations with the different baits showed in Figure 3A (n=2). **B.** Hyperbolic binding curves were fitted on fraction bound values (BI) to each data showed in A to calculate apparent affinities (*K*_app_). 25-30% of the GATA1 protein found in the cell extract was inactive (*f*: active fraction). **C.** The results of the repeated titration experiments with diluted cell extracts (0.1 mg/ml, n=2). Left: 4945_WT bait, right: 4945 haplotype bait. **D.** Validation of measured affinity values and the estimated bait concentrations of the native holdup experiments, by a native holdup titration using a previously characterized palindromic GATA1 bait (n=2) (28). For full blots see Supplementary Figure S3.

Although the nHU assay generates seemingly quantitative data, it has two key limitations that require additional controls. Under the holdup conditions, a large bait excess has to be guaranteed, ensuring that the measured unbound fraction values of the preys are not affected by their total concentrations in the extract. Since DNA baits can interact with a wide range of prey molecules with high apparent affinity, including highly abundant molecules such as histones, there is a possibility of occupying a significant portion of the resin-immobilized bait molecule, causing competition between different preys affecting the outcomes of nHU experiments. To test whether this effect is present, we repeated the nHU titration experiment with the 4945 WT sequence but using 10-fold diluted (0.1 mg/ml total protein concentration) K562 cell extract (Figure 5C, Supplementary Figure S3). No evidence of this inhibitory effect was found, as similar affinities were obtained as before (K_app_=200 nM). Additionally, we use empirically estimated bait concentrations based on prior studies of protein-protein interactions. However, the strong charge of nucleic acids can influence the bait immobilization step, potentially resulting in lower bait concentration estimates than expected, even though we used higher ionic strength during resin immobilization to minimize this effect. To verify our measured affinities, we performed a nHU titration experiment using a palindromic DNA fragment that has been characterized before *in vitro*, using purified recombinant GATA1 protein with surface plasmon resonance (28). In the nHU assay we obtained a dissociation constant (140 nM) very similar to that previously measured with SPR (150 nM), confirming that our bait concentration estimation is in the appropriate range (Figure 5D, Supplementary Figure S3).

### Determination of the effect of disease-associated GATA1 mutations on DNA binding properties by DNA-nHU

We have demonstrated that the nHU approach is suitable to characterize DNA-protein interactions in a quantitative manner. Particularly, nHU titration experiments coupled with Western blot is a quick and straightforward technique to measure the binding of endogenous proteins to various synthetic DNA baits. However, the primary goal of our study was to investigate which pathological GATA1 mutations cause gain or loss of function in DNA binding properties and this requires cell lines that express specific GATA1 mutants. Nevertheless, nHU is an ex-vivo approach where native proteins can be also introduced in the extracts using genetic methods, e.g. by transient transfection. The ectopic expression of GATA1 mutants with a specific tag also reduces reliance on specific antibodies and minimizes issues with antibody recognition due to mutations. For instance, the recognition epitope of the GATA1 antibody used in our experiments is located at the N-terminus of the protein, preventing detection of the GATA1s variant (Supplementary Figure S4). Therefore, we decided to use ectopically expressed HA-tagged GATA1 as prey, as it has been shown before that N-terminal tagging does not affect its DNA binding activity (28).

We used the same HA-GATA1 constructs containing the different mutant versions of the protein than previously in the luciferase assay. We chose HEK293T cell line because it lacks detectable levels of endogenous GATA1 protein or mRNA, preventing any interference with our transiently expressed protein (37). HEK293T cells expressed all selected GATA1 mutants comparably to the wild type protein, and all mutants were detectable with the HA-tag specific antibody (Supplementary Figure S4). As baits in nHU experiments, we selected the WT and the haplotype 51 dsDNA fragments as archetypical tandem, or single GATA motifs (Figure 3A). We performed 12-point nHU titration experiments similarly as shown before and analyzed the results by Western blot against the HA-tag (Figure 6A-B, Supplementary Figure S5-6). Importantly, the ectopically expressed HA-tagged WT GATA1 displayed nearly indistinguishable intrinsic binding affinities towards both tested baits compared to the naturally occurring (untagged) WT GATA1 found in K562 extracts (Figure 5A-B and Figure 6A-B).

**Figure 6.**
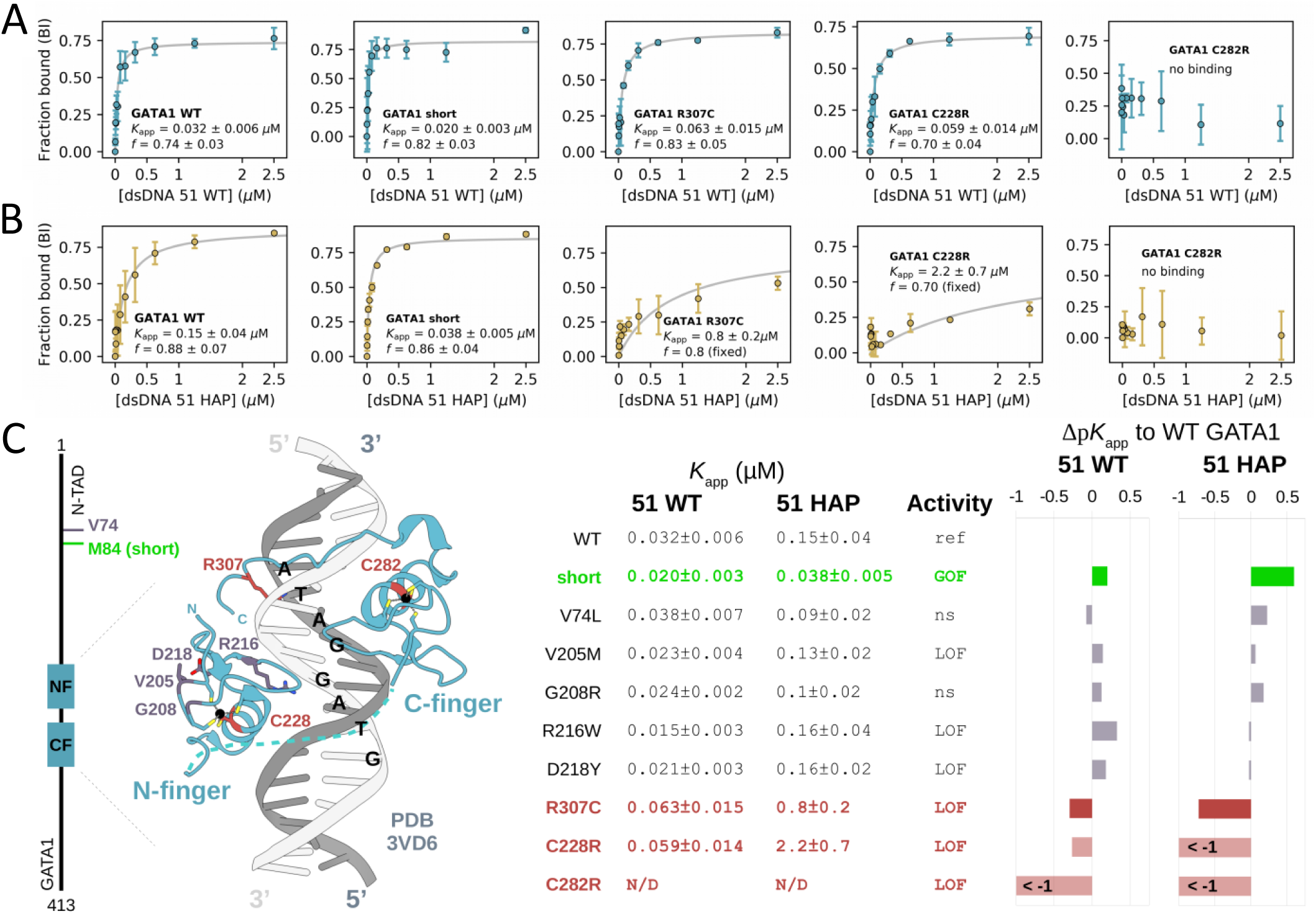
The results of nHU titrations coupled to WB with the different mutant forms of GATA1 using the 51_WT and haplotype sequences and the summary of the effects of different mutations. **A.** The hyperbolic binding equations were fitted to the native holdup titration data and plotted. The 51_WT oligonucleotide was used as archetypal tandem GATA motif. We used HEK293T cell extracts previously transfected with plasmids containing different mutant forms of GATA1 protein. For the Western blot images and the results with other mutants see Supplementary Figure S6. **B.** Hyperbolic binding curves on titrations with the 51_HAP (haplotype) sequence as archetypical single GATA motif (performed as in A). For the Western blot images and the results with other mutants see Supplementary Figure S7. **C.** Summary of the results of the nHU titrations. Left: The different mutations in GATA1 is plotted on the structure of the protein (PDB: 3VD6) (29). Right: *K*_app_ values calculated from native holdup titrations with the 51_WT and 51_HAP sequences. Activity: transcription activation measured in the luciferase assay for the different mutant GATA1 proteins (ref: reference activity (WT), ns: non-significantly different (WT-like), GOF: gain of function, LOF: loss of function). For all blots and fitted plots see Supplementary Figure S5 (51_WT) and S6 (51_HAP).

The artificial mutation affecting the C-terminal zinc finger (CF) resulted in complete loss of activity on both DNA fragments, while the N-terminal zinc finger (NF) mutation reduced the affinity by two-fold for the tandem sequence and approximately 15-fold for the single GATA sequence. This observation is in harmony with previous findings that the NF alone has a very weak affinity for DNA and likely plays a stabilizing and modulatory role after the CF binds to the DNA

(29). Notably, the single GATA site provided much greater differences between the mutant forms of GATA1 compared to the tandem site, which may be due to synergistic effects between neighboring GATA sites. Most pathological GATA1 mutations did not cause major perturbation in DNA binding, including the loss of function V205M, R216W, and D218Y mutations. These data suggest that reduced biological activity originates from other mechanisms than reduced DNA binding. Most importantly, we found that two disease-associated mutations impact DNA binding (Figure 6A-C). The R307C mutation displayed a two-fold decrease in affinity for the tandem GATA motif and approximately a 5.3-fold decrease in affinity for the single GATA motif. In contrast, the short isoform GATA1s exhibited a 1.6-fold increased affinity for the tandem and a four-fold increase for the single GATA site, explaining the increased biological activity observed in the luciferase assay.

It has been suggested before, that the R307C mutation can cause mis-localization, possibly by the disruption of a putative nuclear localization signal (20). To investigate the possibility that mis-localization could play a role in the reduced activity of the protein in the luciferase assay, we examined the co-localization of HA-GATA1 variants with DAPI staining in transfected HEK293T cells. In our assay, all GATA1 variants localized to the nucleus and we did not observe any mis-localization of the different mutant proteins (Figure 7, Supplementary Figure S7). We concluded that the previously observed slight mis-localization might result from the disruption of the DNA-binding capacity of the protein caused by the mutation, and most likely this segment of the protein is not a nuclear localization signal but instead plays a role in DNA binding.

**Figure 7.**
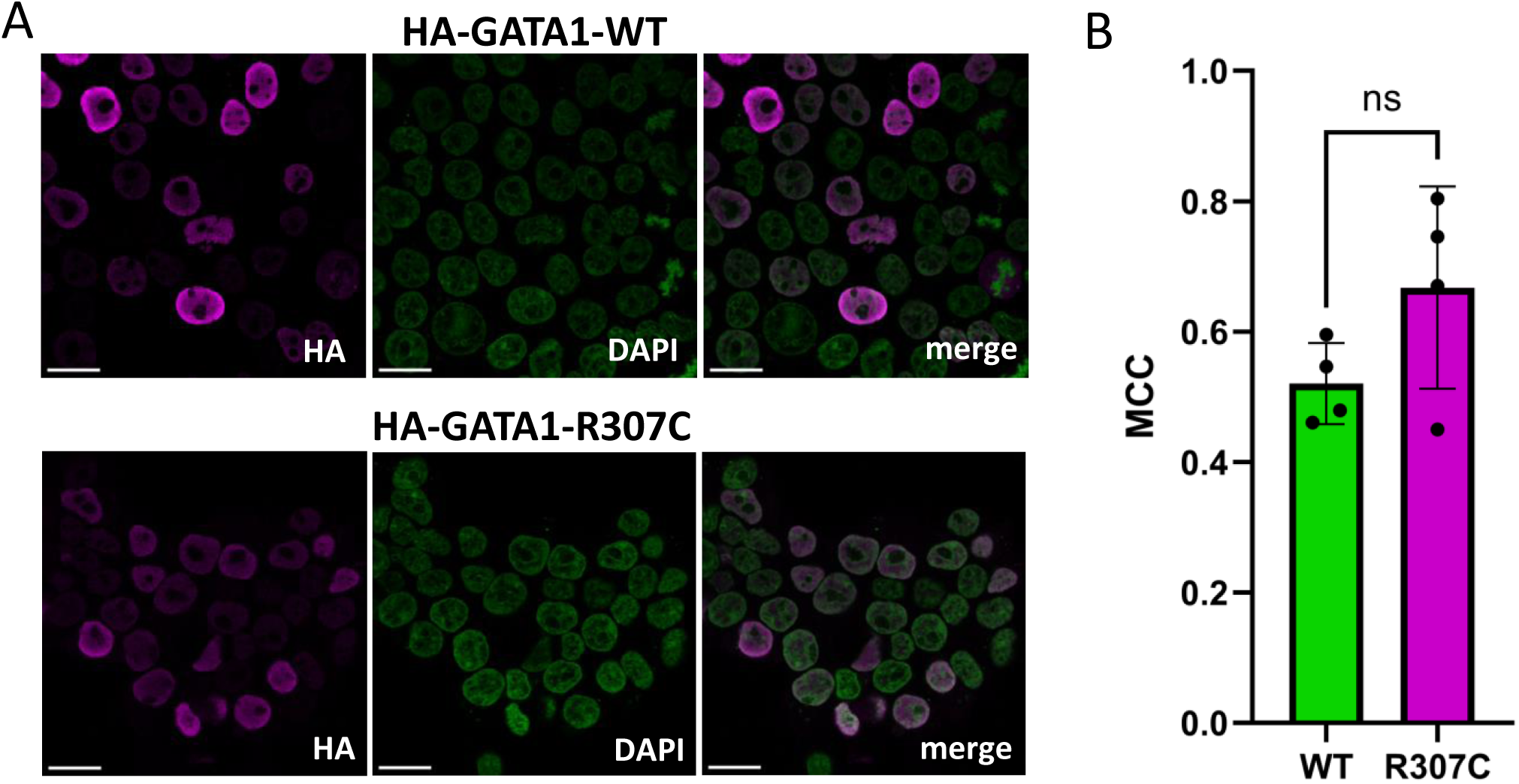
Nuclear localization of the R307C GATA1 protein. **A.** Localization of the HA-tagged WT and R307C mutant GATA1 proteins in transfected HEK293T cells (scale bar: 20 µm). Magenta: HA-GATA1, green: DAPI. For further images see Supplementary Figure S7. B. Co- localization of HA-GATA1 with DAPI was determined by Manders’ correlation coefficient (MCC) tM1 (the fraction of HA-GATA1 in compartments containing DAPI). Statistical significance was determined by two-tailed t-test (ns: non-significant, n=4, on full images excluding dividing cells).

## Discussion

### Functional insight into GATA1 mutations revealed by native holdup

Integrating previous protein-protein interaction data with functional insights from the luciferase assay and binding information from nHU experiments enable us to build a detailed understanding of how disease-associated mutations in GATA1 impact its functions. Out of the four NF mutations investigated here, three (V205M, R216W and R218Y) caused loss of function in the luciferase reporter assay, but we could not measure altered DNA binding for these variants (Figure 6C). These mutations structurally localize to the same surface of the protein, and most likely disrupt the binding of an important co-activator, such as FOG1, LMO2 or TAL1, leading to reduced transcriptional activation by GATA1 indirectly, as suggested before (6, 11, 30, 33). Of note, one of the studies also found that mutations of these positions caused different degrees of loss of function, due to the differences in structural incompatibility of the different GATA1 protein-protein interaction interfaces (30).

In contrast to the NF mutations, two natural variants directly affected DNA binding: the R307C variant and the short isoform. The R307C variant causes a decrease in affinity which manifests in a reduced cellular activity. This aligns well with the artificial zinc-finger mutations that abolish the DNA binding, causing a complete loss of function. It has previously been suggested that mutation in this position may abolish a nuclear localization signal, but our results indicate that it is more likely that the DNA-binding capacity of this mutant is impaired, which may cause the slight mis-localization observed before (20). Indeed, structurally, it has been previously described that this arginine is located in the extension of the CF, protruding deep into the minor groove, and most probably contributes to the DNA binding (29) (Figure 6C). It has also been proposed that phosphorylation at Ser310 may influence the protein’s function, which is disrupted by the R307 mutation (53). However, further analysis is needed to understand the interplay between the protein’s phosphorylation states and the DNA-binding affinity.

In contrast, increased DNA binding affinity was observed for the short GATA1 variant, which also showed enhanced cellular activity in the luciferase assay. GATA1s was observed before in DS- TAM and DS-AMKL, and that the expression levels of the short isoform correlated with the severity of TAM (54). These data suggested that this mutant may disrupt the delicate balance of regulation by GATA1 leading to cellular overproduction of precursors in the trisomy 21 genetic background (25–27). The observed allosteric effect on DNA binding can be explained by the removal of a potential inhibitory control in the N-TAD of GATA1. This mutation has been shown to specifically disrupt chromatin occupancy and H3K27 methylation pattern at erythroid-specific genes and lineage specific effects on maintaining progenitors (21, 23, 55), indicating that, despite the stronger binding observed in this study, it may affect the transcriptional activity differently across various genomic sites. Overall, by integrating previous protein-protein interaction results with comparative functional activity data and a systematic comparison of DNA binding activities measured by nHU, our study provides a comprehensive understanding of the molecular mechanisms underlying the disease-associated mutations in GATA1. This approach not only elucidates the specific alterations in DNA binding affinity caused by these mutations but also offers valuable insight into how these changes contribute to the diverse cellular phenotypes observed in GATA1-related diseases, such as severe anemia, thrombocytopenia, Down syndrome-related transient abnormal myelopoiesis (DS-TAM), and acute megakaryocytic leukemia (DS-AMKL).

### Opening new frontiers in transcription factor research with native holdup

In this study, we aimed to characterize previously described pathological mutations in GATA1 by investigating different aspects of TF activity. While existing methods can systematically assess the quantitative effects of mutations on promoter activation, a significant gap remains in methodologies for studying DNA-protein interactions. To address this, we have developed the present native holdup assay, which enables the quantitative measurement of interactions between a synthetic or purified DNA bait and a full-length prey directly taken from cell extract. Using the erythroid-specific promoter segments of the *ATP2B4* gene as a model system, we investigated the DNA binding properties of GATA1 mutants transiently expressed in mammalian cells. Our results demonstrate that this versatile method can quantitatively measure DNA binding affinities of TFs without requiring protein purification, making this an ideal approach for systematically studying TF mutations.

DNA-binding proteins play a critical role in regulating gene expression by interacting with specific regions in the genome. Mutations in these proteins, particularly those associated with disease, can disrupt their ability to bind DNA effectively, leading to altered transcriptional regulation and contributing to pathogenesis. Despite their importance, many DNA-binding proteins remain poorly characterized, and we lack comprehensive knowledge of how disease-associated mutations impact their interactions with DNA. While computational approaches, including innovative AI tools like AlphaFold and AlphaMissense, excel at predicting structure-disrupting mutations, they fall short in assessing more subtle changes, such as mutations that slightly alter DNA-binding affinity or shift DNA-binding profiles (56, 57).

Experimentally, researchers have traditionally used pulldown-based assays to identify DNA-protein interactions. However, the dynamic nature of many interactions means that the partners may dissociate during the washing steps, leading to incomplete or misleading results. Our native holdup (nHU) assay addresses these challenges by enabling the identification of DNA-binding partners without the need for washing steps, making this method ideal for capturing even transient interactions. This method offers a robust and simple way to systematically investigate the effects of disease-associated mutations in either the DNA or in the DNA-binding protein. The nHU assay can not only identify the partner of a given DNA bait, but also provides a quantitative measurement of intrinsic affinity. Such data are typically difficult to obtain and require the production of purified proteins at high quantities. These are not requirements of a nHU assay that can be done with affordable synthetic DNA oligos, a small amount of commercially available resin and simple cell extracts. The results that these assays provide are comparable to conventional biophysical assays, such as ITC or SPR, or other methods routinely used in the nucleic acid fields, such as mobility shift assays. In the past, multiple studies measured DNA binding to GATA1 using quantitative biophysical approaches, yet only with a few exceptions these studies were performed with isolated zinc fingers of GATA1, containing no or mild, non-disruptive mutations, often at non-physiological conditions (29, 30). In the case of nHU, there is no need to purify partner proteins, hence even highly unstable, yet biologically important full-length proteins that would be not suitable for conventional methods can be studied at near native conditions. In conclusion, nHU offers new perspectives for studying nucleic acid-protein interaction and due to the simplicity of the assay, it can be effortlessly implemented in any cell biology laboratory.

## Acknowledgements

We thank the staff of the IGBMC Cell Culture and Light Microscopy Facility for their help in cell culturing and imaging.

## Author contributions

B.Z. conceptualized the study, designed and performed most of the experiments. B.Z. and G.G. designed and performed all the nHU and did all the final data analyses. B.M. and L.N. performed the MS experiments and primary data analysis. O.M. and B.S. took part in conceptualization and helped in the luciferase assay. G.T. obtained funding for the experiments. G.T. and G.G. supervised the project. B.Z. and G.G. wrote the original draft, and all authors reviewed the manuscript.

## Supplementary Figure Legends

**Supplementary Figure S1.** Western blot images on transfected K562 cell extracts that were used in luciferase assay, to determine the HA-GATA1 expressions. **A.** Western blot images of replicates. **B.** Statistical analysis of expressions of different GATA1 mutants. *: p<0.05 with two-tailed t-test.

**Supplementary Figure S2.** Western blot images of single point nHU experiments with 51_WT, SCR1 and SCR2 sequences.

**Supplementary Figure S3.** Western blot images of titration nHU experiments with the 4945_WT, 4945_HAP, 51_WT, 51_HAP, or with the CTRL_PALINDROME_ sequences on K562 cell extracts either at 1 mg/ml total protein concentration (upper panel) or 0.1 mg/ml total protein concentration (lower panel).

**Supplementary Figure S4.** Validation of transient expressions of the different mutant versions of GATA1 in HEK293T cells by Western blot. The HA antibody recognizes all mutants, but the GATA1 antibody doesn’t recognize the short version of GATA1 (GATA1s) as the epitope recognized by the antibody might located at the N-terminal part of the protein.

**Supplementary Figure S5. A.** Western blot images of nHU titrations with the 51_WT sequence on cell extracts prepared of HA-GATA1 WT and mutant transfected HEK293T cells. **B.** Determined hyperbolic binding curves on titrations with the 51_WT bait and with the mutant GATA1 proteins that are not shown in Figure 6.

**Supplementary Figure S6. A.** Western blot images of nHU titrations with the 51_HAP sequence on cell extracts prepared of HA-GATA1 WT and mutant transfected HEK293T cells. **B.** Determined hyperbolic binding curves on titrations with the 51_HAP bait and with the mutant GATA1 proteins that are not shown in Figure 6.

**Supplementary Figure S7.** Additional images on the localization determination of different mutant versions of the GATA1. HEK293T cells were transfected with HA-GATA1 WT or mutant plasmids and immunostaining was carried out using an antibody recognizing the HA epitope (magenta), and the nucleus was visualized by using DAPI staining (green). Scale bar: 20 µm.

## Supplementary Data Legends

**Supplementary Data 1.** Results of the luciferase assay measurements and the related Western blots for expression determination. The first sheet contains the luminescence values measured with transfected K562 cells and an *ATP2B4* erythroid-promoter construct. The assay was performed with 3 biological and 2 technical replicates, data were normalized that the average of HA was 0 and the WT was 100 (%). The second sheet contains expression result by densitometry on Western blots, on the same samples as the luminescence was measured. For the Western blot images see Supplementary Figure S3.

**Supplementary Data 2.** Results of the nHU-MS experiments. The first four sheet contains the measured log_2_ fold change values, statistical significances (-log_10_P), calculated binding intensities, apparent affinities and significance thresholds. In cases where the significance threshold is smaller than the obtained statistical significance of the measurement, the apparent affinity is considered significant and a numerical affinity value is displayed in the p*K*_d_sign column. Proteins that are classified as transcription factors are highlighted in the last column. The last tab contains an assembled affinity matrix of all significant affinities.

**Supplementary Data 3.** Results of the GO enrichment. On the first sheet, an assembled affinity matrix of all significant affinities is shown with the last column showing the number of significant measurements for each prey. GO enrichment analysis was performed for all proteins with >0 values in this column or for the promiscuous proteins for proteins with a value of 4 in this column. The following sheets contain the significant GO terms for cellular compartments or for molecular functions.

**Supplementary Data 4.** Results of the nHU-MS experiments of the scrambled DNA fragments. Similarly to Data 2, each sheet contains the measured log_2_ fold change values, statistical significances (-log_10_P), calculated binding intensities, apparent affinities and significance thresholds and the significant affinities. The first three sheet contains the measurement of the SCR1, SCR2 and the 51WT baits against a biotin control. The last two sheets contain the measurement of the 51WT bait against the SCR1, or SCR2 controls.

