## Supplemental figures for "Quantitative analysis of DNA-GATA1 binding alterations linked to hematopoietic disorders"

### Supplementary Figure S1

A

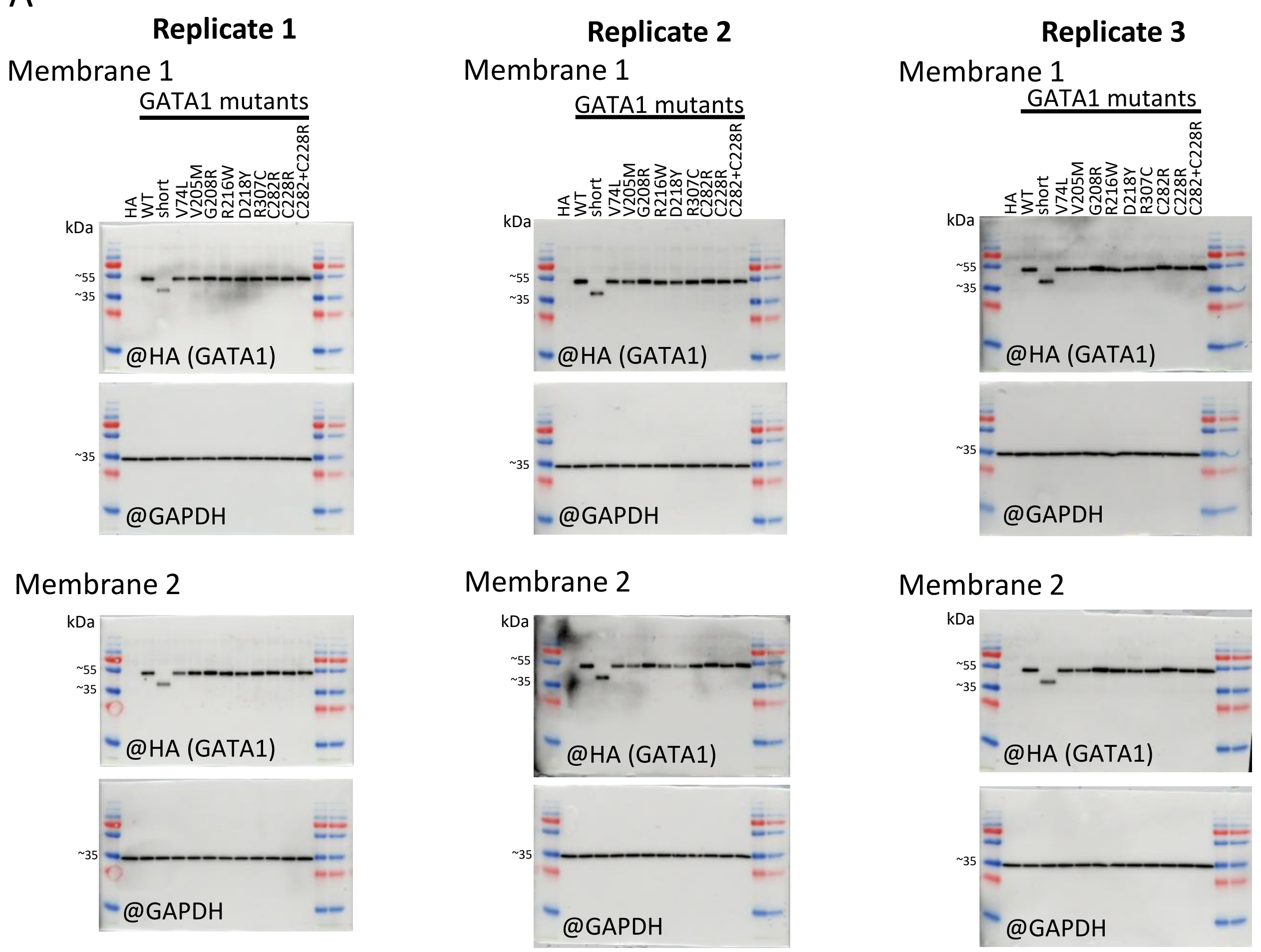


B

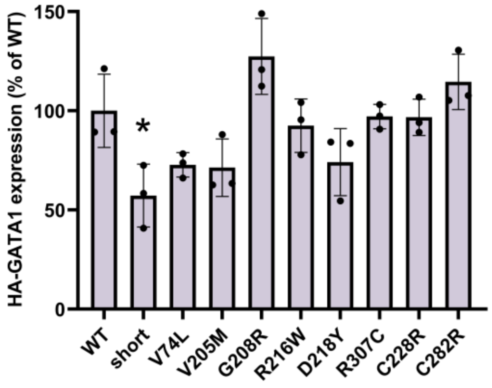

### Supplementary Figure S2

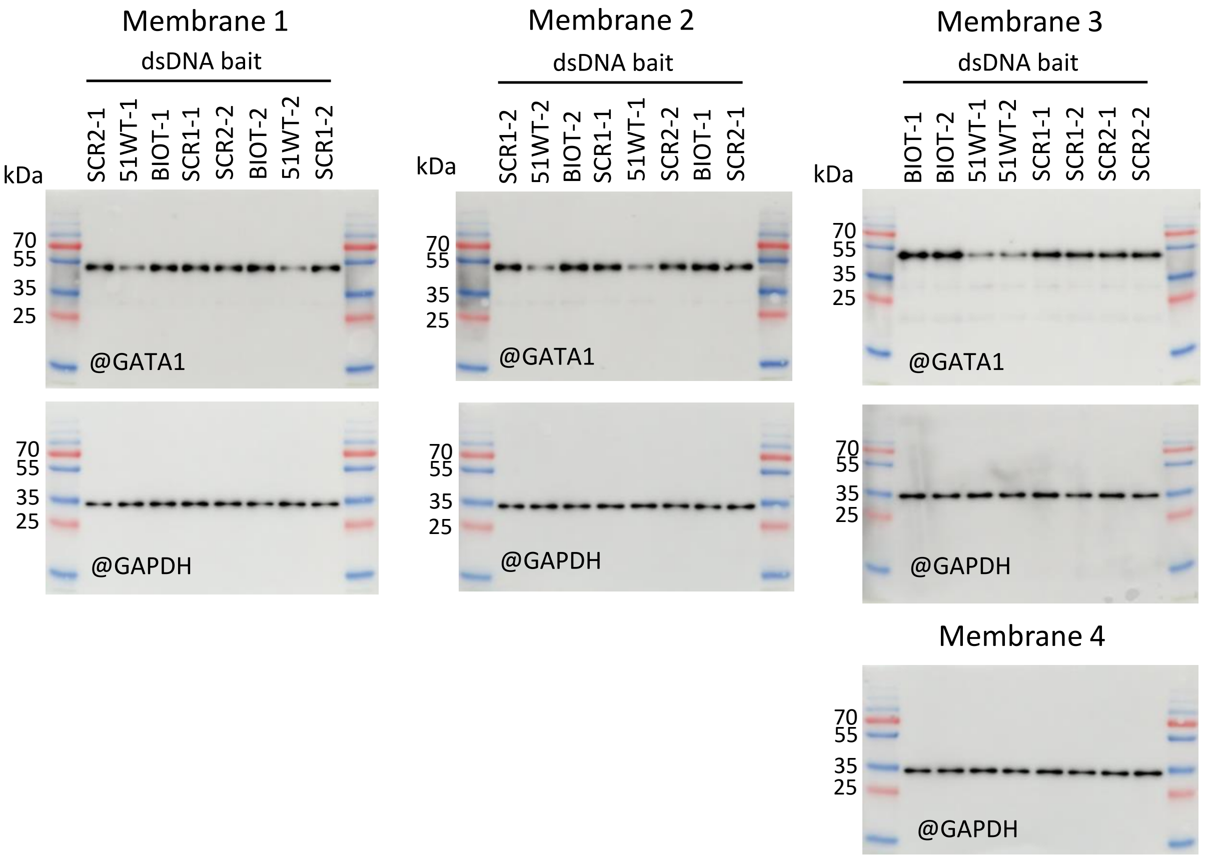

### Supplementary Figure S3

**Experiments @ 1 mg/ml total protein concentration**

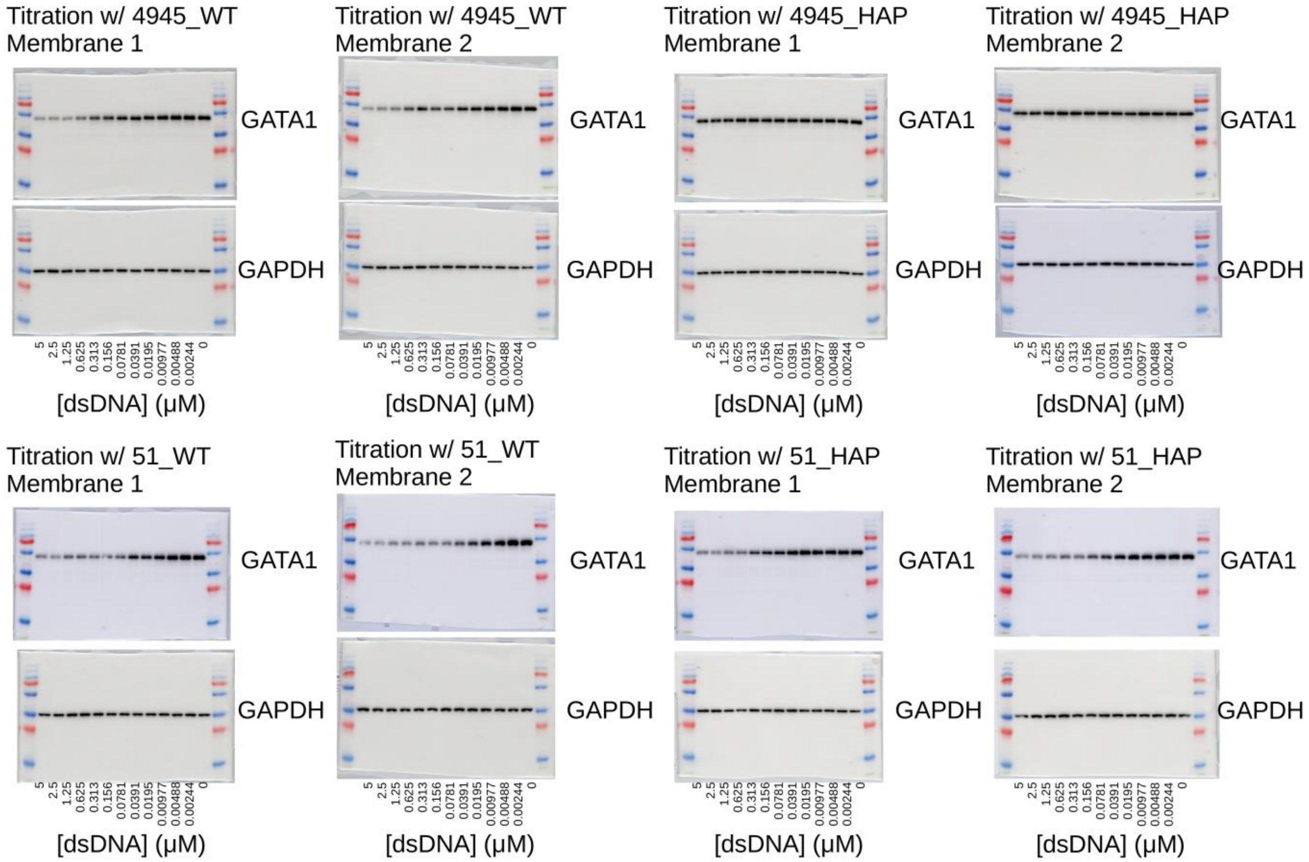


**Experiments @ 0.1 mg/ml total protein concentration**

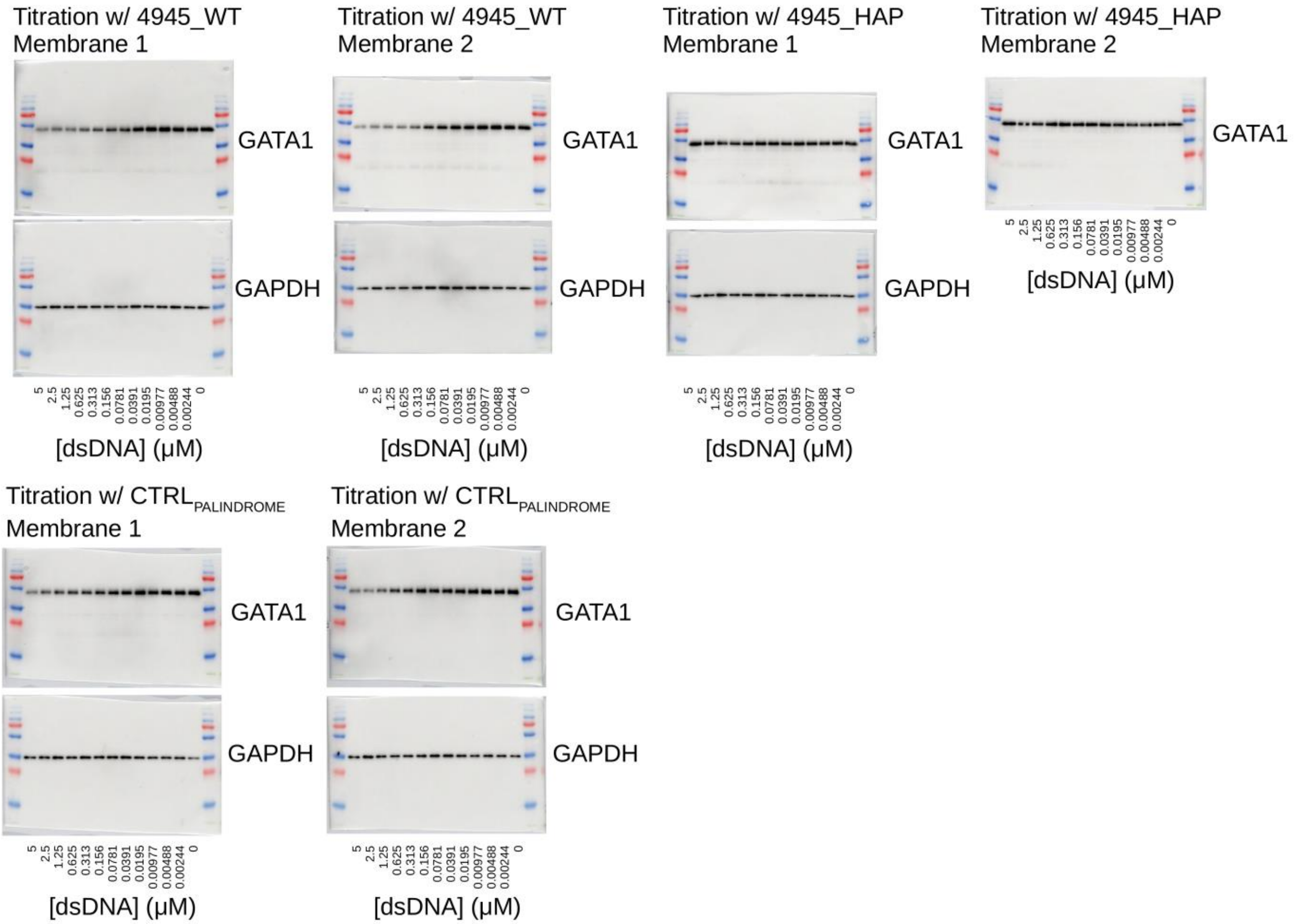

Supplementary Figure S4

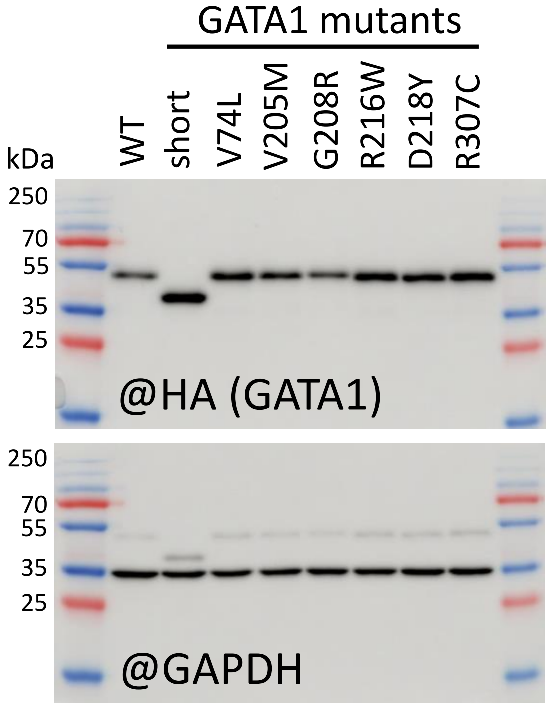


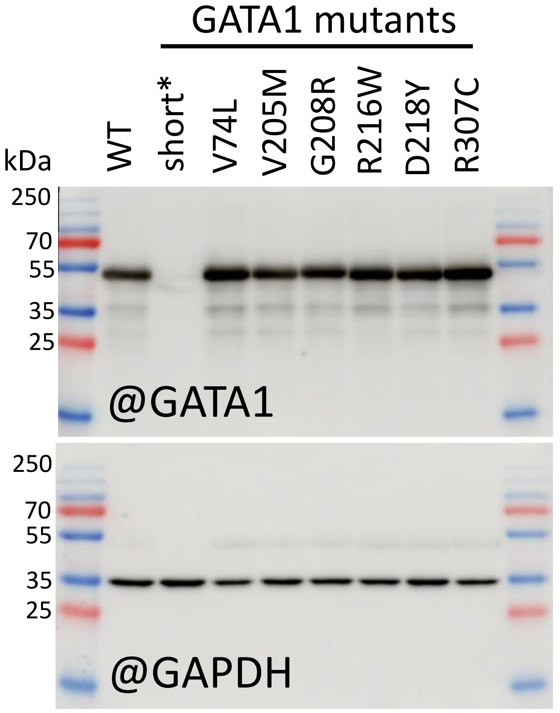


* antibody epitope probably in the N-TAD

### Supplementary Figure S5

A

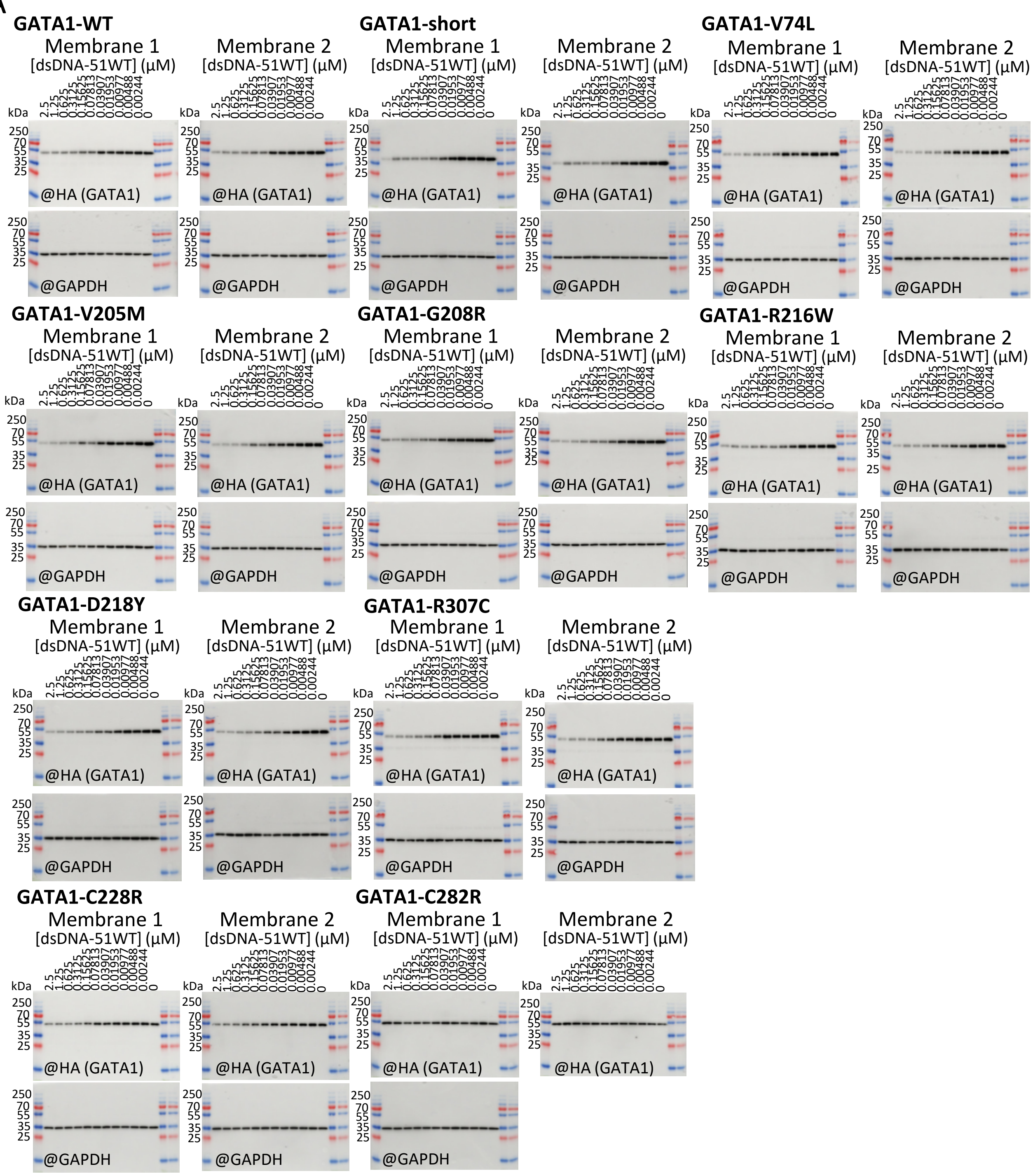


B

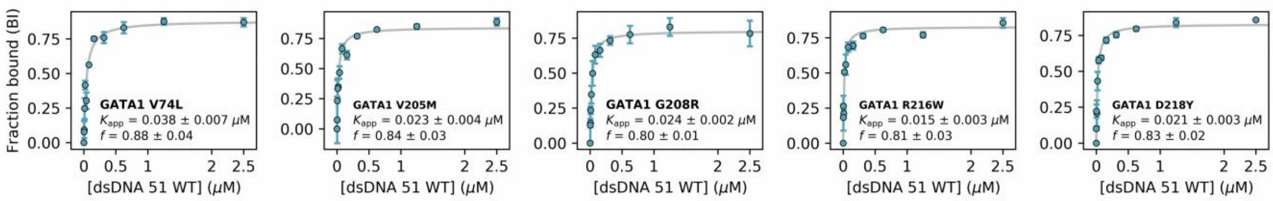

### Supplementary Figure S6

A

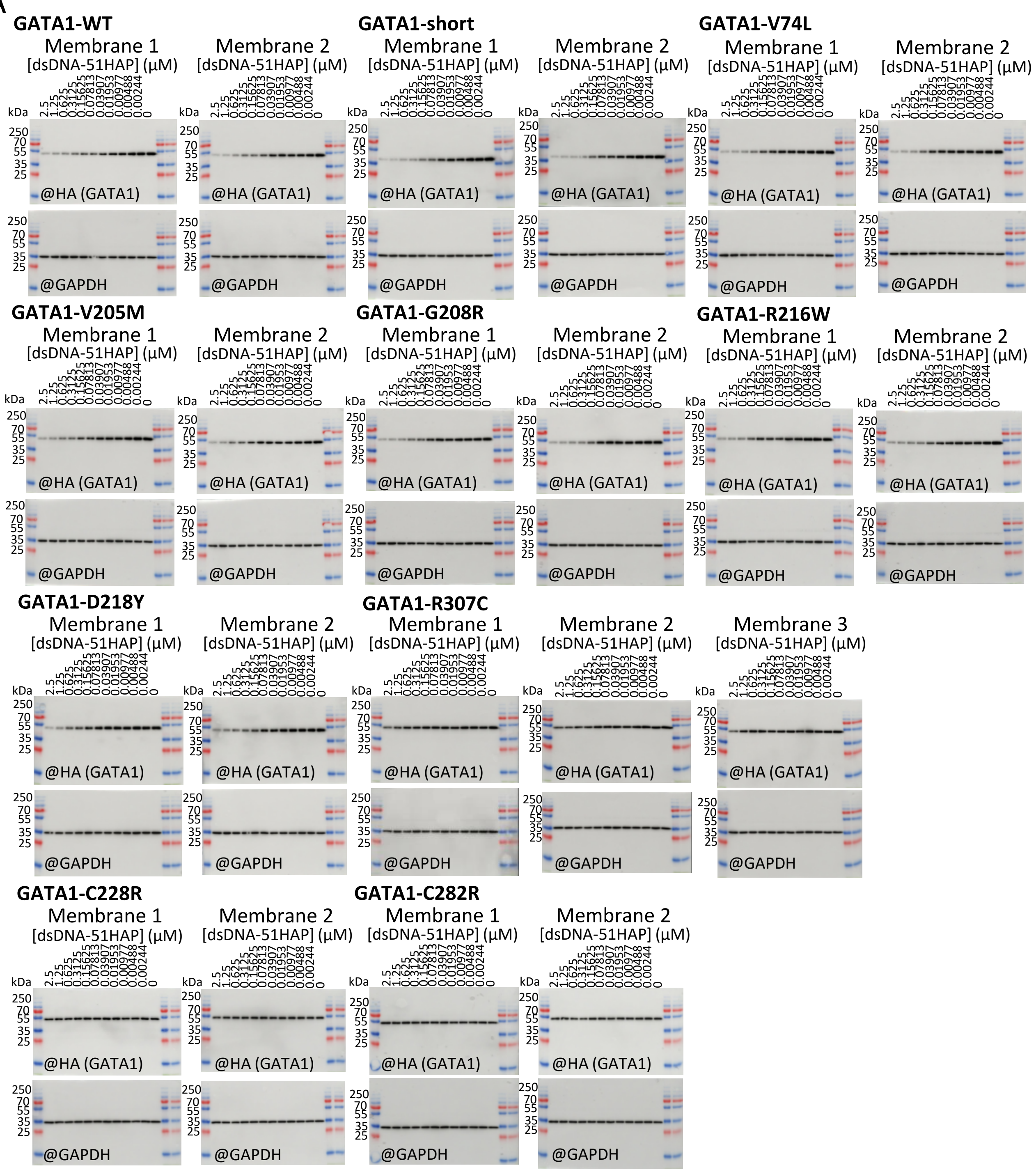


B

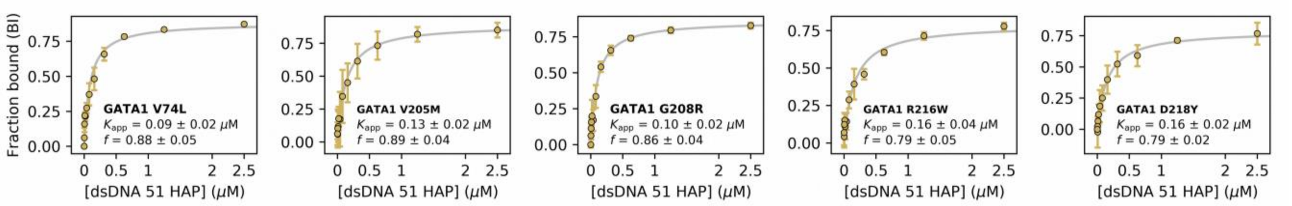

Supplementary Figure S7

A

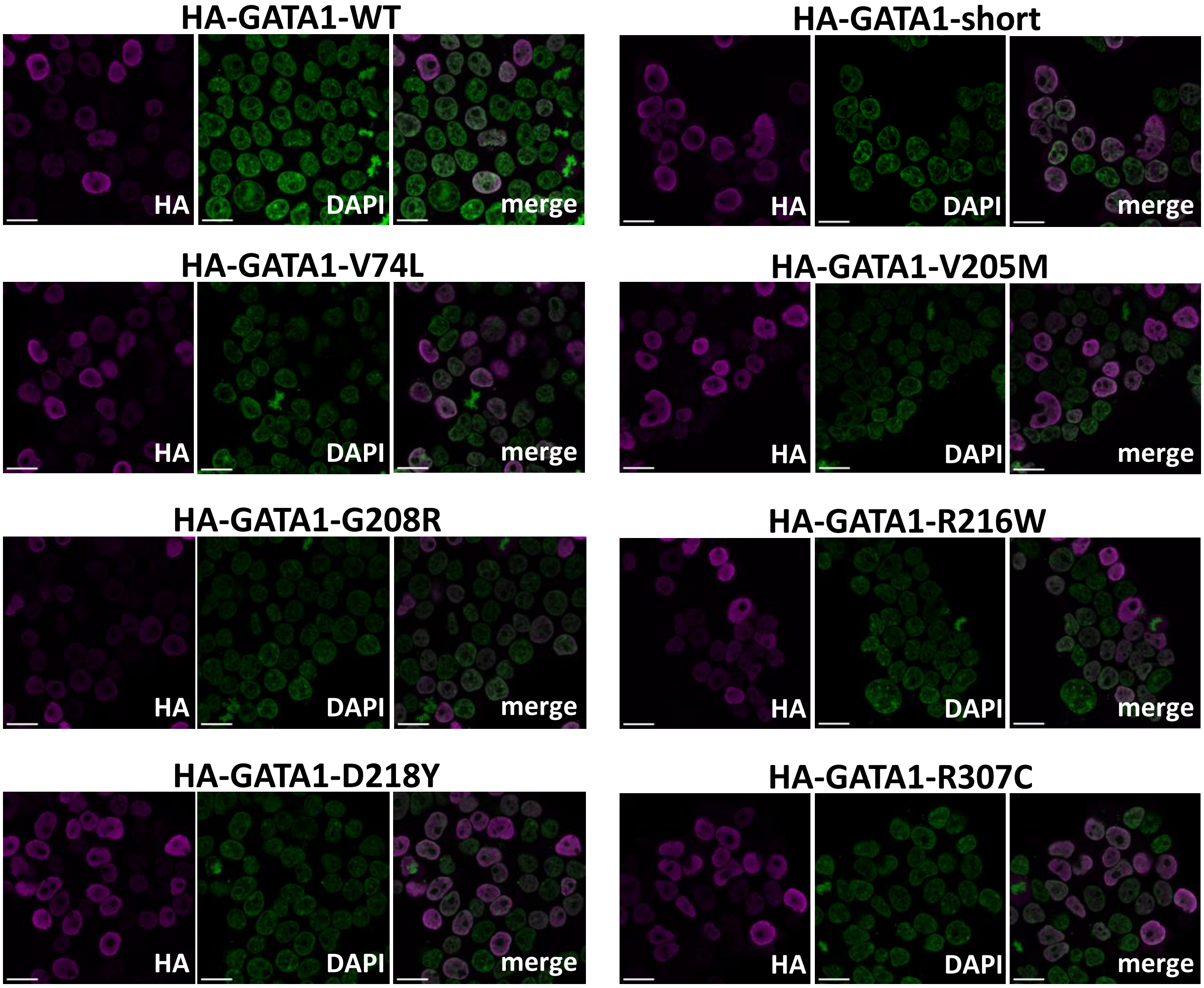
